## Supplementary Information for "VIA: Generalized and scalable trajectory inference in single-cell omics data"

**Shobana V. Stassen[1], Gwinky G. K. Yip[1], Kenneth K. Y. Wong[1,3], Joshua W. K. Ho[2,4] and Kevin K. Tsia[1,3]**
[1]Department of Electrical & Electronic Engineering, The University of Hong Kong, Pokfulam Road, Hong Kong
[2]School of Biomedical Sciences, Li Ka Shing Faculty of Medicine, The University of Hong Kong, Pokfulam, Hong Kong
[3]Advanced Biomedical Instrumentation Centre, Hong Kong Science Park, Shatin, New Territories, Hong Kong
[4]Laboratory of Data Discovery for Health, Hong Kong Science Park, Shatin, New Territories, Hong Kong

### Table of Contents

1. Table of benchmarked datasets (Table S1)
2. Performance characteristics of benchmarked methods (Table S2)
3. Runtime summary of benchmarked methods (Table S3)
4. Key VIA parameters (Note S1)
5. Parameter Settings for benchmarked methods (Table S4-S5)
6. Derivation of Expected Hitting Time for lazy-teleporting random walk (Note S2)
7. Summary of F1-Score Lineage Detection 'Stress-Tests' for datasets and methods (Fig. S1)
8. Topological Benchmarking on Synthetic Datasets (S2-S5) (Note S3)
9. Benchmarking on real datasets (Fig. S6-S26) with gene-trend and noise analyses (Note S4)
10. VIA performance directly on genes without PCA: Synthetic and real data (Fig. S27-S29) (Note S5)
11. VIA performance with Kmeans in lieu of PARC (Fig. S30-S32) (Note S6)
12. Detailed explanation of VIA's MOCA graph (Note S7)
13. FACED image-based biophysical features and the corresponding definitions (Table S6-S7)

### 1. Table of Datasets Benchmarked

| | Single cell modality | Organism | Number of cells | Time series/ snapshot | Biological Process | Reference |
|---|---|---|---|---|---|---|
| **1. Pre-B** | scRNA-seq | Mouse | 500cells x 9,075 genes | time series 2-24 hours | pre-BI to pre-B2 | Gomez-Cabrero et al., 2019 |
| **2. Human embryoid** | scRNA-seq | Human | 16,825 ESC | time series (5 intervals 0 - 27 days) | Embryogenesis | Moon et al., 2019 |
| **3. Hematopoeisis** | scRNA-seq | Human | 5500 cells x ~15,000genes | snapshot | Hematopoeisis | Setty et al., 2019 |
| **4. MOCA** | scRNA-seq | Mouse | 1.3 Million after filtering | time series (E9.5-E13.5) 5 intervals | Organogenesis | Cao et al., 2019 |
| **5. Pancreatic** | scRNA-seq | Mouse | 2531 cells x ~ 15,000 genes | snapshot | Endocrine dev | Bastidas-Ponce 2019 |
| **6. mESC mesoderm** | CyTOF | Mouse | 90,000 cells x 28 antibodies | time series (daily for 11 days) | ESC to Mesoderm cells | Ko et al., 2020 |
| **7. Cell cycle** | Imaging | Human | ~4000 | snapshot | cell cycle (2 breast cancer types ) | in-house (present work) |
| **8. Hematopoesis** | scATAC-seq | Human | 2034 x 8192 TF | snapshot | Hematopoesis | Buenrostro et al. 2018 |
| **9. Cardiac progenitor** | scATAC + scRNA-seq | Mouse | 695 cells+ 197 cells | snapshot | cardiac progenitor lineage | Jia et al. 2018 |
| **10 a - i. Toy topologies** | Simulated | Simulated | 1000 cells x 1000 genes | | | Saelens et al. DynTOY |

**Table S1 Summary of datasets employed in the present study.**

### 2. Performance characteristics of benchmarked methods

| Method | Scalability | Multi-omic | Cell Fate Prediction | Pseudotime | Complex Topology | Lineage paths |
|---|---|---|---|---|---|---|
| VIA | High | Yes | Yes | Yes | Yes | Yes |
| PAGA | High | Yes | No | Yes (DPT) | Yes | No |
| Palantir | Medium | Yes | Yes | Yes | No | Yes |
| Slingshot | Low | Yes | Yes | No | No | Yes |
| CellRank | Low | No | Yes | No | No | Yes |
| Monocle3 | Medium | No | Yes | Yes | Yes | Yes |
| STREAM | Medium | Yes | Yes | Yes | No | Yes/No |

**Table S2 General comparison of the state-of-the-art TI methods.** Many state-of-the-art TI methods can automate lineage related analysis. However, based on our extensive performance testing, they generally face challenges in meeting the demands of increasingly large, diverse (multi-omic) and complex single-cell data. They are:

**Scalability: (maximum capacity of cells processed within 10 mins)**

- High: able to process 200,000 cells <10 mins
- Medium: able to process 20,000 cells <10 mins
- Low: able to process 2000 <10 mins

**Multi-omic:** Accepts (in theory) any single-cell feature set as input (not limited to scRNA-seq)

**Cell Fate prediction:** Predicts cell fates corresponding to differentiated or temporally evolved cell types

**Pseudotime:** Computes unified ordering of cells that can be used downstream to compare lineages

**Complex Topology:** Captures cyclic, disconnected trajectories (or hybrids thereof) as well as trees

**Lineage paths:** Computes pathways most likely to represent differentiation along each lineage

### 3. Runtime summary of benchmarked methods

| Datasets | Cells | Dimensions after pre-processing | VIA | Palantir | PAGA | Slingshot | CellRank | Monocle3 | STREAM |
|---|---|---|---|---|---|---|---|---|---|
| MOCA | 1,300,000 | 30 PCs | 40 | 270 | 180 | X | X | X | 390 |
| Mesoderm ESC | 89,782 | 28 proteins | 2 | 6 | 10 | 390 | NA | NA | 15 |
| Endocrine | 2,531 | 30 PCs | 1 | 3 | 2 | 3 | 10 | 10 | 1 |
| Human CD34+ | 5500 | 200 PCs | 2 | 4 | 3 | 150 | NA | 15 | 2 |

**Table S3 Runtime (minutes) for benchmarked methods.** Filtering steps, visualization of results (e.g. UMAP) and pre-processing (e.g. PCA) are excluded. 'X' denotes either a memory error or prohibitive runtimes. Monocle3 completed the initial projection and low-dimensional clustering in ~90 mins but encountered a memory error in the graph learning step (exceeding 120GB). Slingshot took 390 mins on the 90K mESCs and thus deemed unfeasible for MOCA. In the case of CellRank, spliced and unspliced count matrices were only available for the endocrine data on which it took 10 mins to parse 2500 cells - the bottleneck was the scVelo step required to infer directionality using the spliced/ unspliced counts.

### 4. Supplementary Note 1: Key parameters in VIA

As an exploratory TI tool, VIA provides a transparent interface that allows the user a great deal of control in reconfiguring parameters. That being said, all tests reported in this paper ran VIA on its default parameters. We also performed multiple stress tests on each of the diverse datasets considered to show that the default parameters are sensible, and that changes to key algorithmic and input parameters consistently produce reliable results. We briefly comment on some parameters in VIA and the reasonable ranges within which they can be tuned.

**PARC clustering parameters (Step 1 Fig 1):** VIA calls upon PARC in its default state, but the number of K (nearest neighbors), level of pruning, and resolution parameter can all be adjusted (refer to PARC[1.]). For all datasets except MOCA (in consideration of runtime), VIA was run in two iterations, the first being coarse-grained (corresponds to PARC default) as this is best for terminal state inferences, and the second where PARC forces over-fragmentation in the clustering stage (by reducing the allowed size of clusters as a percentage of overall population) to get a more fine-grained resolution of the pseudotimes but retaining the terminal-states from the initial iteration.

**VIA laziness and teleportation:** For all runs presented, we have set laziness to 5% and teleportation to 1%. Reasonable ranges for laziness are 1-10% and for teleportation: 1-5%

**VIA graph pruning for visualization of topology:** For visualization purposes, the cluster-graph can be pruned (either by retaining the top *n* edges, where *n* is user defined, or by thresholding using a statistical threshold) to provide a clearer illustration of the structure - this does not impact the inferred underlying trajectory or related characteristics such as pseudotime or terminal state prediction.

**Imputation:** VIA includes a gene-imputation option using the pre-computed HNSW graph. This is only used (optionally) for computing gene-trends, and not for trajectory inference. The power to which the diffusion operator is powered can be modified (similar to MAGIC[2], the default value is 3).

### 5. Parameter setting for benchmarked methods

| Process | Type | Dimensions | Starting Point | Range of parameters varied | Remarks |
|---|---|---|---|---|---|
| **Synthetic** | DynToy | 1000 cells x 1000 features | M1 (Milestone 1) | K-NN=20, PC=10 for composite accuracy | Each composite score comprises 5 metrics and thus we fixed the number of KNN and PCs for calculating the composute accuracy of synthetic data |
| **Hematopoeisis** | scRNA-seq | 5500 cells x 14651 genes | HSC | K-NN={10,20,...80}<br>PC ={10,20,...,200} | Both knn and PC are varied to conduct a more comprehensive gridsearch |
| **Hematopoeisis** | scATAC-seq | 2034 x 8192 TF | HSC | K-NN=20<br>PC = {10,20,...,200} | We consider the results for two different pre-processing pipelines and limit the parameter range to only varying the PCs |
| **Pancreatic** | scRNA-seq | 2531 x 10,000 ?? genes | Endocrine Progenitor | HVG = {300,500,1000,2000,...,10000}<br>PC = {30,40,...,80} | Here we primarily consider the effect of varying the number of HVGs |
| **MOCA** | scRNA-seq | 1.3 Million Cells | E9.5 Epithelial cell | K-NN={20,30,...,50} , PC = {30}<br>PC = {20,30,..,50}, KNN = {30} | We visually compare outputs of methods that are able to process this dataset within 3 hours |
| **Cardiac** | scRNA-seq/ scATAC-seq | 695 cells+ 197 cells | Cardiac Progenitor | K-NN=10 (knn=15 for Palantir which otherwise fails at lower knn=10) | Here we primarily consider the effect of varying the number of PCs. K (KNN) is set to a low value since the the cell count is small |
| **mESC mesoderm** | CyTOF | 90,000 cells x 28 antibodies | Day 0 cell | All features used w/o PCA, except Slingshot which could not handle the full-feature set at high cell counts<br>K-NN **= {10,20,30,...,}** | Sample time annotations allow us to compute pseudotime correlations for this dataset for different K values using all 28 antibodies directly (except Slingshot which required PCA to handle the high cell count) |
| **Cell cycle** | Imaging | 4000 cells x 38 morphological features | G1 cell/ G1 cluster | All features are used<br>K-NN=20 | The 38 features (without PCA) are run directly. Subsets of these features (e.g. excluding volume related features) are probed |

**Table S4: Summary of key parameters considered in the benchmarked datasets.** In particular, the table also summarizes the choice of root (i.e. starting point of the trajectory) and the rationale for which parameters are probed to impact TI for each dataset.

| Method | General Parameter settings with emphasis on why/when parameters are changed from default | Starting point selection |
|---|---|---|
| **VIA** | • Underlying PARC settings were left at default graph pruning and Leiden resolution.<br>• We showed that good results are achievable using other clustering such as kmeans (n_clusters set to 20 as a default) which shows that the VIA method is not overly dependent or sensitive to clustering methods, as long as a sufficient resolution is captured. | Provided as a single cell index corresponding to an early cell for real datasets; provided as a group-level label for more data-driven starting point selection for Synthetic data where Milestone group annotations are available |
| **PAGA** | • Set the number of KNN for both the PCA and diffusion map iteration<br>• The number of PCs and diffusion components in both knn-graph stages is set to the number of PCs selected rather than allowing the diffusion components to be subset to the default 10 DCs<br>• To plot gene trends we need to manually select appropriate terminal clusters and then manually select a corresponding set of clusters that make up a lineage-pathway | A single cell index correpsonding to a suitable early cell (the same starting cell is given to all methods requiring a single-cell root) |
| **Slingshot** | • KNN is not a parameter in Slingshot and hence not 'grid searched'.<br>• We first tested the method using the recommended GMM clustering but found this to be too coarse to detect the relevant lineages. The other option was using KMeans where we typically set K=15 to allow for slight overclustering without adding too much fragmentation. Slingshot's visualization of lineages is linked to the input PCs which makes it difficult to visualize results on more complex data where 2 PCs are not a good basis for visualization. | Root manually selected as the cluster containing HSCs and the 'start' of the trajectory |
| **Monocle3** | • Monocle3 requires UMAP to be applied to the PCs before computing TI. This is recommended on their tutorial pages as part of their intended pipeline and also prompted by the tool when running it. As such, the KNN is set as the number of neighbors used in the UMAP computations and the PC value is the number of top PCs used as UMAP input.<br>• The default distance metric set by Monocle in using UMAP is "cosine distance" but this works very poorly for the datasets at hand and we therefore switched to Euclidean distance which significantly improved results | Root branch is manually selected after getting the branching structure and selecting the most suitable milestone based on the ground truth annotations of early cells (progenitors, stem cells) |
| **STREAM** | • For STREAM, the original use case suggests to run PCA followed by one of UMAP, MLLE or Spectral Embedding (SE). To ensure consistency in benchmarking and reduce the impact of additional feature reduction and subsetting incurred by a second stage of dimensionality reduction, we chose to only perform the PCA step and skip further dimensionality reduction (a more detailed explanation is given below).<br>• SE was generally too coarse for complex real data (often yielding a simple bifurcation) and MLLE tends to overfragment the data such that almost each cell type (early, intermediate and late) corresponds to a "leaf branch". MLLE was thus sensitive the number of components retained and quickly reached a breakpoint where overfragmentation occured. UMAP, like SE, was often too coarse and limited the granularity.<br>• Due to the uncertainty introduced by choice of secondary Dimension reduction (with no clear winner), we choose to apply only PCA and skip the second dimensionality reduction step. This also makes for a fairer comparison to palantir, paga, Slingshot and VIA which are computed on the PCs.<br>• For the Cardiac scATAC-seq /scRNA-seq datasets, Stream would not run on the PCs (i.e. produced errors) so we used SE after PCA.<br>• We follow the tutorials provided by STREAM to incorporate a two stage approach suggested for more complex data. This allows you to refine the graph by adding more branches and nodes in a second iteration.<br>• All other parameters such as the number of initial clusters, the incremental clusters used in intializing and seeding the graph, were left as default (as per the tutorial).<br>• KNN is not a relevant parameter unless SE/UMAP/MLLE are used. | Root branch is manually selected after getting the branching structure and selecting the most suitable branch based on the ground truth annotations of early cells (progenitors, stem cells) |
| **Palantir** | • Set the number of PCs and KNN in run_diffusion_maps() and run_pca() function and override the default PC and KNN settings which otherwise subset the DCs to 10.<br>• The number of sampling points is left at the default 1500 as Setty et al., 2019 show robustness to this parameter | A single cell index correpsonding to a suitable early cell (the same starting cell is given to all methods requiring a single-cell root) |
| **CellRank** | Ran as shown on the tutorial page to reproduce the results shown in their paper for endocrine data. Skipped for other datasets as they did not have unspliced counts | Automatically determines the starting point. Since this would sometimes incur spurious starting points, we compared gene trends of the pancreatic dataset for an output which detects the correct root. |

**Table S5: Summary of settings selected for the benchmarked methods.** Method-specific parameter settings and rationales for non-default choices are provided for each of the benchmarked methods.

#### 6. Supplementary Note 2: Derivation of Closed-Form Formula for Expected Hitting Time of Lazy, Teleporting, Random Walk

The cluster graph constructed in VIA is mathematically defined as a weighted connected graph **G** ($V$, $E$, $W$) with a vertex set $V$ of $n$ vertices (or nodes), i.e. $V = \{v_1, \cdots, v_n\}$ and an edge set $E$, i.e. a set of ordered pairs of distinct nodes. $W$ is an $n \times n$ weight matrix that describes a set of edge weights between node $i$ and $j$, $w_{ij} \geq 0$ are assigned to the edges $\left(v_i, v_j\right)$. For an undirected graph, $w_{ij} = w_{ji}$. Let $P$ be the probability transition matrix ($n \times n$) of a standard random walk on this graph G can be given by

$$P = D^{-1}W \tag{1}$$

where $D$ is the $n \times n$ degree matrix, which is a diagonal matrix of the weighted sum of the degree of each node, i.e. the matrix elements are expressed as

$$d_{ij} = \begin{cases} \sum_k w_{ik} & ,i = j \\ 0 & ,i \neq j \end{cases} \tag{2}$$

where $k$ are the neighbouring nodes connected to node $i$. Hence, $d_{ii}$ (which can be reduced as $d_i$) is the degree of node $i$.

The Laplacian $L$, and the normalized Laplacian $L_{NL}$ are defined as follows. The $i^{th}$ eigenvector of the normalized Laplacian is denoted by $\Phi_i$ and its corresponding eigenvalue $\eta_i$.

$$L = D - W \tag{3}$$
$$L_{NL} = D^{-0.5} L D^{-0.5} \tag{4}$$

where $\Phi_1 = \bar{1}$ and $\eta_1 = 0$. Using Eqs. (3) and (4), we also make note of the relationship between the probability transition matrix $P$ (Eq. (1)) and $L_{NL}$ which we will use later on:

$$L_{NL} = D^{-0.5}(D - W)D^{-0.5} = I - D^{-0.5} W D^{-0.5} \tag{5}$$
$$I - P = I - D^{-1}W = L_{NL} = D^{-0.5}(I - D^{-0.5} W D^{-0.5}) D^{0.5} = D^{-0.5} L_{NL} D^{0.5} \tag{6}$$

The $\varphi-$normalized Laplacian $L_\varphi = \varphi I + L_{NL}$ has the same eigenvectors as $L_{NL}$, and eigenvalues $\varphi + \eta_i$. F. Chung and S. Yau[3.] show that for $\varphi-$normalized Laplacian, the modified Green's function $R_\varphi$, where $L_\varphi R_\varphi = I$, can be written as:

$$R_\varphi = \sum_{i=1}^{n} \frac{\Phi_i \Phi_i^T}{\varphi + \eta_i} \tag{7}$$

We note that the Green's function is a symmetrized form of the personalized PageRank[4.] $pr_\alpha$. In order to compute the expected hitting time $h_\alpha(q, r)$ from node $q$ to node $r$ in terms of the Green's function, we first express $h_\alpha(q, r)$ in term of $pr_\alpha$.

$$h_\alpha(q, r) = \frac{\left[pr_\alpha(e_r)^T\right](r)}{d_r} - \frac{\left[pr_\alpha(e_r)^T\right](q)}{d_q} \tag{8}$$

where $0 < \alpha < 1$ and where $e_i$ is an indicator vector with 1 in the $i^{th}$ entry and 0 elsewhere (i.e. $s_m = 1$ if $m = i$ and $s_m = 0$ if $m \neq i$).

In order to describe a lazy and teleporting random walk and its hitting times, we need to formulate the Green's function of a linearly scaled and shifted Laplacian. We denote this generalized Laplacian, $L_{\varphi,NL} = \varphi I + kL_{NL}$, which has the same eigenvectors as $L_{NL}$, but eigenvalues equal to $\varphi + k\eta_i$. The generalized Green's function of the generalized Laplacian $L_{\varphi,NL}$ can be written as

$$R_{\varphi,NL} = \sum_{m=1} \frac{\Phi_m \Phi_m^T}{[\varphi + k\eta_m]} \quad (9)$$

$$L_{\varphi,NL} R_{\varphi,NL} = I \quad (10)$$

We now construct a *lazy-teleporting* random walk, Z, and extract the corresponding Laplacian $L_{\varphi,NL}$. Z has probability ( $1 - x$ ) of being lazy (where $0 < x < 1$ ), i.e. staying at the same node, then

$$Z = xP + (1 - x)I \quad (11)$$

where $I$ is the identity matrix. When teleportation occurs with a probability ( $1 - \alpha$ ), the modified lazy-teleporting random walk $Z'$ can be written as

$$Z' = \alpha Z + (1 - \alpha)\frac{1}{n}J \quad (12)$$

where $J$ is an $n \times n$ matrix of ones. The personalized PageRank vector $pr_\alpha(s)$ (defined as a column vector here) is the unique solution to

$$pr_\alpha(s)^T = \alpha pr_\alpha(s)^T Z + (1 - \alpha)s^T \quad (13)$$

Where $s$ is the seed vector of the initial probability distribution across the $n$ nodes (such that $\sum_m s_m = 1$, where $s_m$ is the probability of starting at node $m$)[37,4]. For ease of manipulation, we write $\beta = \frac{2(1-\alpha)}{(2-\alpha)}$.

Re-writing the pagerank equation (Eq. (13)) in terms of the transition matrix $P$ and collecting like terms, we can extract the Laplacian for the modified random walk as follows:

$$pr_\alpha(s)^T = \alpha pr_\alpha(s)^T Z + (1 - \alpha)s^T$$
$$\Rightarrow \alpha pr_\alpha(s)^T [xP + (1 - x)I] + (1 - \alpha)s^T$$
$$\Rightarrow pr_\alpha(s)^T 2\frac{(1-\beta)}{(2-\beta)}[xP + I - xI] + \frac{\beta}{(2-\beta)}s^T$$

Collecting $pr_\alpha(s)^T$, we obtain

$$(2I - \beta I)pr_\alpha(s)^T = pr_\alpha(s)^T [2xP - 2\beta xP + 2(1 - x)I - 2\beta(1 - x)I] + \beta s^T$$
$$\Rightarrow pr_\alpha(s)^T [\beta I - 2xP + 2\beta xP + 2xI - 2\beta xI] = \beta s^T$$
$$\Rightarrow pr_\alpha(s)^T [I(2x - 2\beta x) - P(2x - 2\beta x) + \beta I] = \beta s^T$$

Using the relation $I - P = D^{-0.5} L_{NL} D^{0.5}$ (Eq. (6)), we extract the scaled and shifted expression for the modified Laplacian $L_{\beta,NL}$ corresponding to the lazy-teleporting random walk:

$$pr_{\alpha}(s)^{T} \left[D^{-0.5}(L_{NL}(2x - 2\beta x) + \beta I) D^{0.5}\right] = \beta s^{T} \tag{14}$$

Since $R_{\beta,NL}$ is the inverse of the modified Laplacian, we can rearrange Eq. (14) as

$$pr_{\alpha}(s)^{T} = \beta s^{T} D^{-0.5} R_{\beta,NL} D^{0.5} \tag{15}$$

We substitute $\varphi = \beta$ and $k = 2x(1 - \beta)$ in Eq. (9) and obtain:

$$R_{\beta,NL} = \sum_{m=1} \frac{\Phi_m \Phi_m^T}{[\beta + 2x(1-\beta)\eta_m]} \tag{16}$$

Substituting Eq. (15) into our expression for expected hitting time Eq. (8), we can write the hitting time in terms of the closed form expression of $R_{\beta,NL}$. Making use of the fact that $\frac{1}{d_r} = \left[D^{-1} e_r\right](r)$, and $D^{-0.5} R_{\beta,NL} D^{-0.5}$ is symmetric such that[37]

$$\chi_r^T [D^{-0.5} R_{\beta,NL} D^{-0.5}] \chi_q = [D^{-0.5} R_{\beta,NL} D^{-0.5}]_{rq} = \chi_q^T [D^{-0.5} R_{\beta,NL} D^{-0.5}] \chi_r, \tag{17}$$

we can express the expected hitting time as

$$h_{\alpha}(q, r) = \beta (e_r - e_q)^T D^{-0.5} R_{\beta,NL} D^{-0.5} e_r \tag{18}$$

### 7. Summary of F1-Score Lineage Detection ‘Stress-Tests’ for multiple datasets and methods

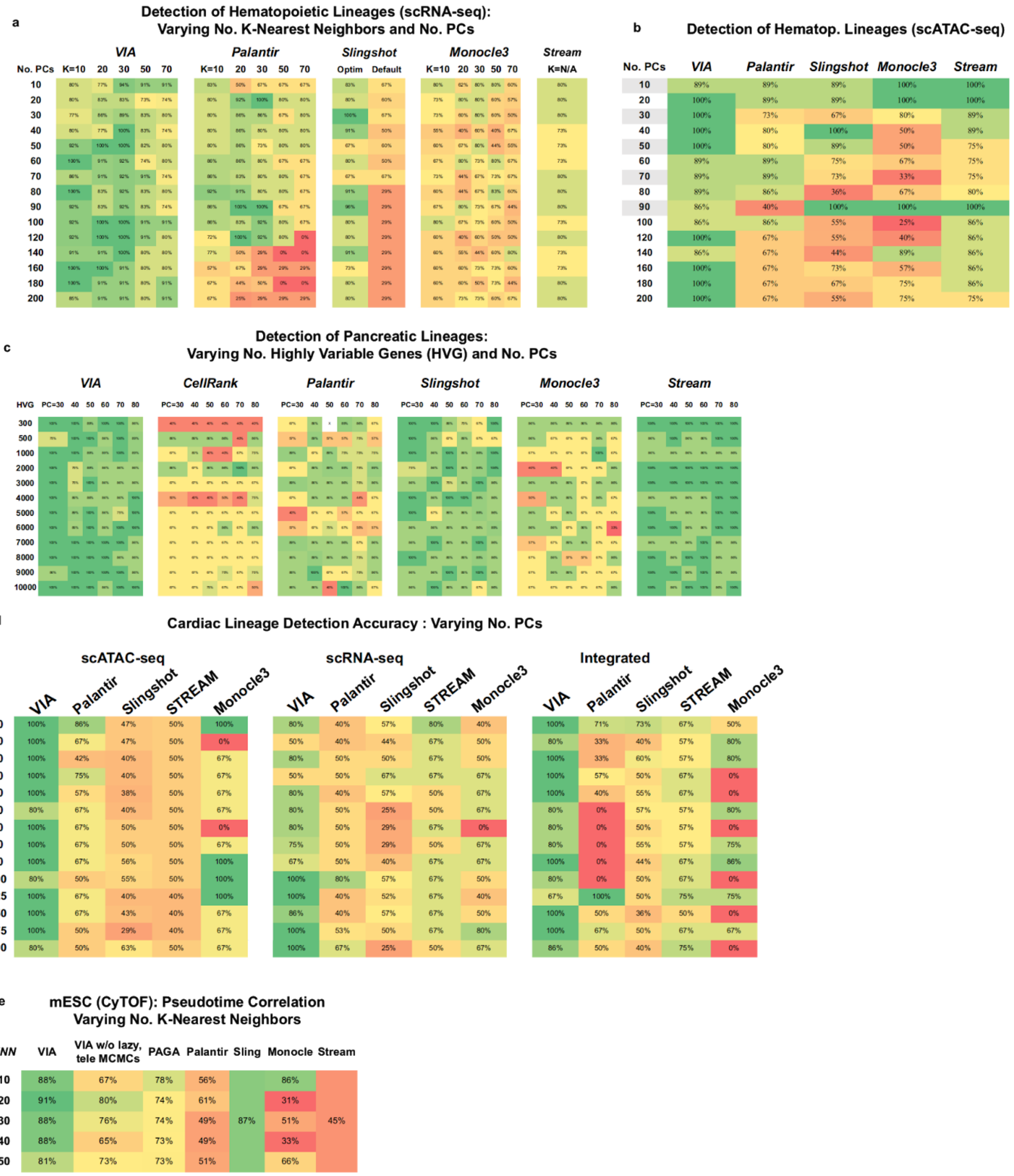


**Fig.S1 Reconfigurability of VIA.** Summary of various parameter stress-tests on datasets benchmarked with other TI methods where possible.

### 8. Supplementary Note 3: Benchmarking on synthetic data with complex topologies

We use 9 synthetic datasets to benchmark the performance of VIA and other methods on a variety of trajectories and topologies. The 9 synthetic datasets are generated using DynToy and contain 1000-3000 cells across 1000 features. They span (bi/multi) furcations, cyclic, connected (which are combinations of cyclic and furcating) and disconnected (which are combinations of all the aforementioned). The composite accuracy metric assesses multiple layers of the inferred trajectory, taking into account the topological similarity between the reference model and the inferred topology, the correlation between the real and 'pseudo' times, as the prediction accuracy of the terminal cell fates (lineages). For certain metrics, the absolute measurements of similarities or differences are converted into a percentage scale such that 100% corresponds to the highest score. The composite metric is the arithmetic mean of the 5 scaled metrics.

**Ipsen-Mikhailov[39] (IM)**: IM distance is used to measure the similarity of global graph topology. The IM ranges from 0 to 1 and equals the difference in spectral densities of two graphs. $IM(G_1, G_2) = 0$ implies that the graphs are isospectral. Unlike the Graph Edit Distance and the F1-branch score, where only the structure of each link's immediate neighbourhood contributes to the distance value, the IM considers the structure of the whole topology. We use $1 - IM$ for the composite accuracy score.

$$IM(G_{TI}, G_{REF}) = \sqrt{\int_0^{\infty} [\rho_{REF}(\omega) - \rho_{TI}(\omega)]^2 d\omega} \tag{19}$$

$$\rho(\omega) = \sum_{k=1}^{N-1} \frac{\gamma}{(\omega - \omega_k)^2 + \gamma^2} \tag{20}$$

$\rho(\omega)$ is the graph spectral density as a sum of Lorentz distributions, $\omega_k$ are the vibrational frequencies given by $\sqrt{\lambda_i}$ where $\lambda_i$ are the eigenvalues of the Graph Laplacian. $\gamma$ is the half width at half maximum of the Lorentz distributions.

**Graph Edit Distance (GED)**: is defined as the cost of converting $G_{TI}$ to $G_{REF}$ with the least possible number of operations. Each operation has a cost of one and includes insertion/deletion of edges *and* nodes. Oftentimes graph edit distances like the (e.g. GED, Hamming distance) are normalized by the entire set of possible edges $N(N - 1)$, but this creates an inflated sense of accuracy in the case of very sparse graphs like the reference topologies of the synthetic datasets. We therefore normalize by the sum of nodes and edges in $G_{REF}$. 1- 'Normalized' GED is used in the composite accuracy measure.

**F1-Branch score:** is applied to the local branch accuracy and defined as follows. A False Negative edge in the inferred model is when there is an edge in the reference model connecting clusters/cell types/groups that is absent in the inferred trajectory. A False Positive edge in the inferred model is when an edge occurs in the inferred model that is not actually present in the reference model. A True Positive Edge is one that exists in the inferred model that also exists in the reference. These three measurements are used to compute the F1-score.

$$F_1 = \frac{tp}{tp + 0.5(fp+fn)} \tag{21}$$

**Temporal Correlation:** Given the sampling times of the synthetic data, we use the Pearson Correlation coefficient as a measure of how closely the inferred pseudotime follows the true sampling times. $\sigma_X$ is the standard deviation of X, and $\mu_X$ is the mean.

$$\rho_{x,y} = \frac{E[(X-\mu_X)-(Y-\mu_Y)]}{\sigma_X \sigma_Y} \tag{22}$$

**F1-Cell Fate score:** We use the harmonic mean of recall and precision as given by Eq. (21) to quantify the prediction accuracy of terminal states. *tp* is a true-positive: the identification of a terminal cluster that is in fact a final differentiated cell fate; *fp* is a false positive identification of a cluster as terminal when in fact it represents an intermediate state; and *fn* is a false negative where a known cell fate fails to be identified.

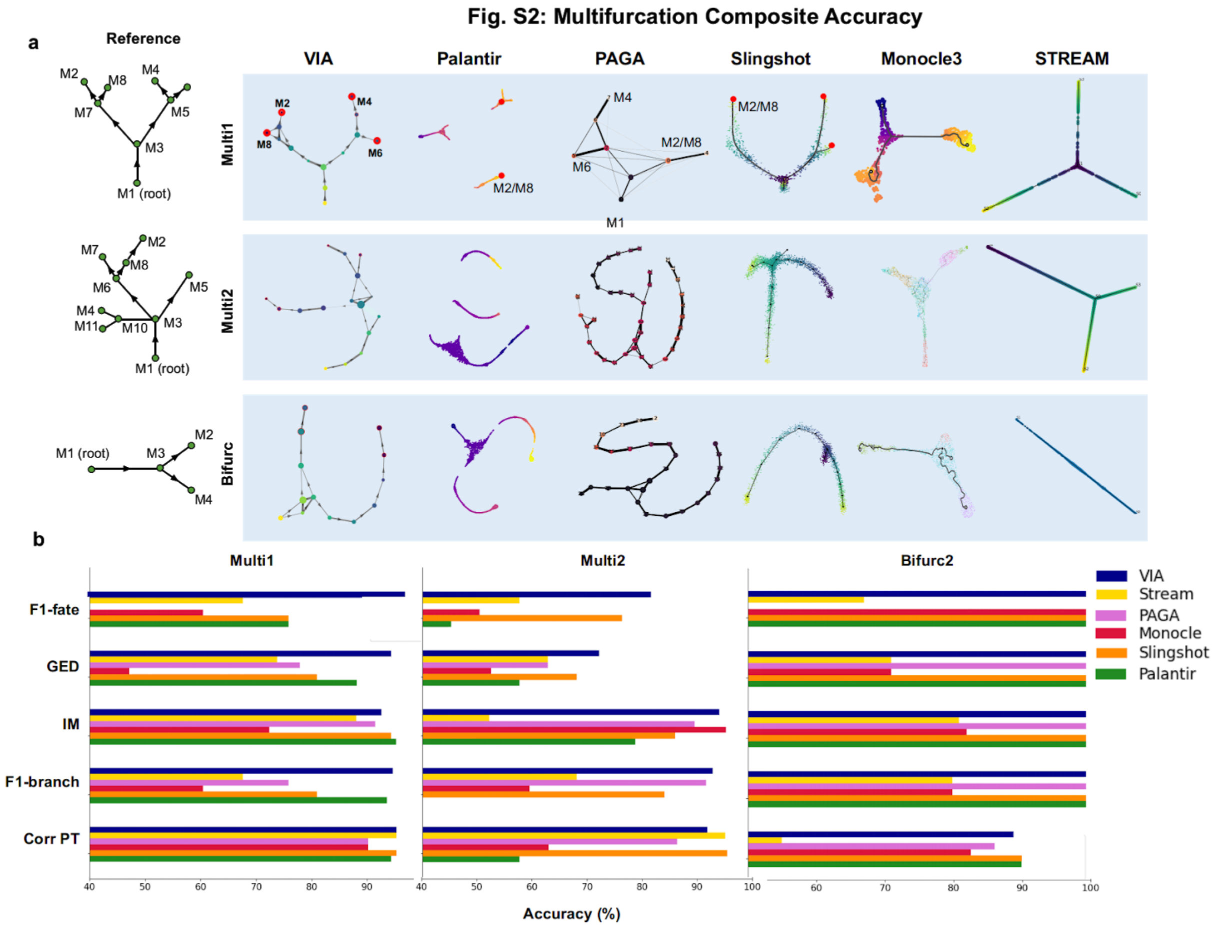


**Fig. S2 Benchmarking on multifurcating synthetic datasets. (a)** Sample outputs by each TI method shown for each of the multifurcating datasets colored by their inferred pseudotimes. The reference topologies are shown on the left. **(b)** The accuracy scores of each of the 5 individual metrics (F1-cell fate score (F1-fate), GED, IM, F1-branch and pseudotime correlation (Corr PT)) comprising the composite accuracy score are shown individually.

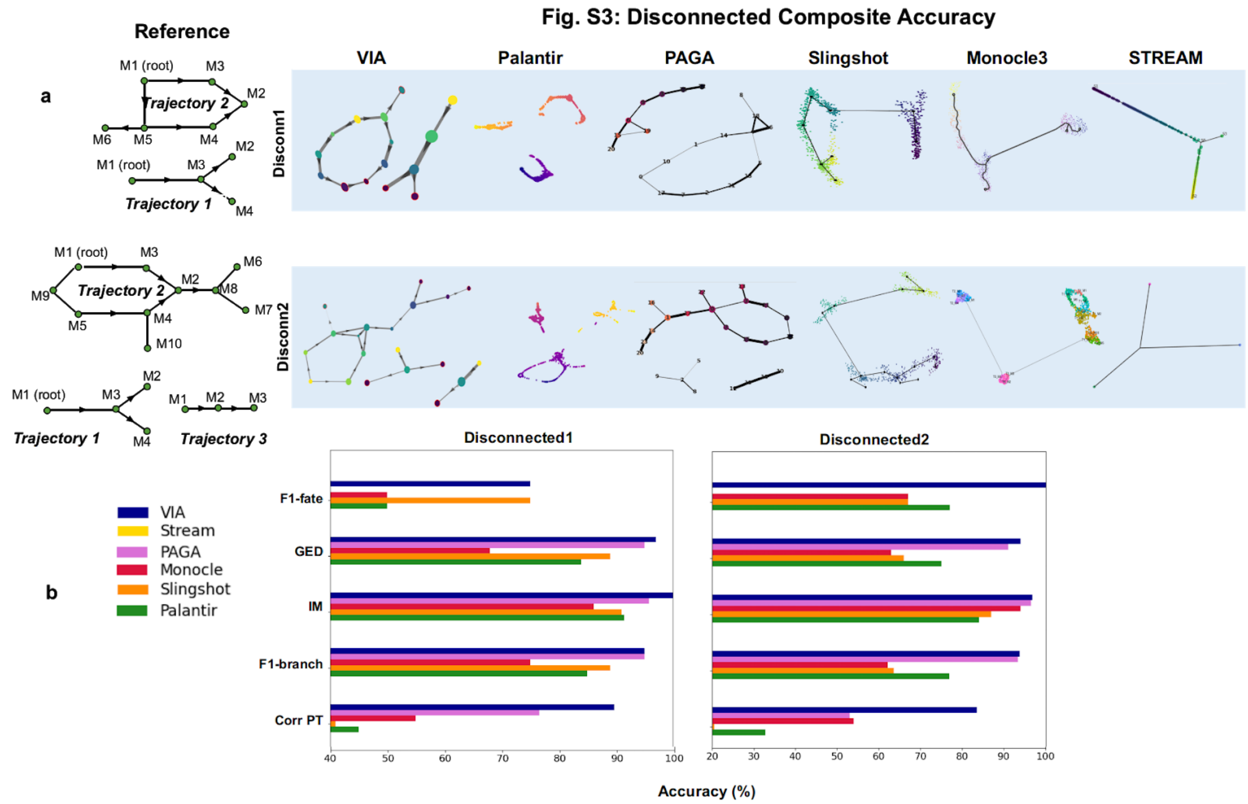


**Fig. S3 Benchmarking on disconnected synthetic datasets. (a)** Sample outputs by each TI method shown for the two disconnected datasets, each comprising a combination of sub-trajectories as shown in the reference topologies (left). **(b)** The accuracy scores of each of the 5 individual metrics (F1-cell fate score (F1-fate), GED, IM, F1-branch and pseudotime correlation (Corr PT)) comprising the composite accuracy score are shown individually.

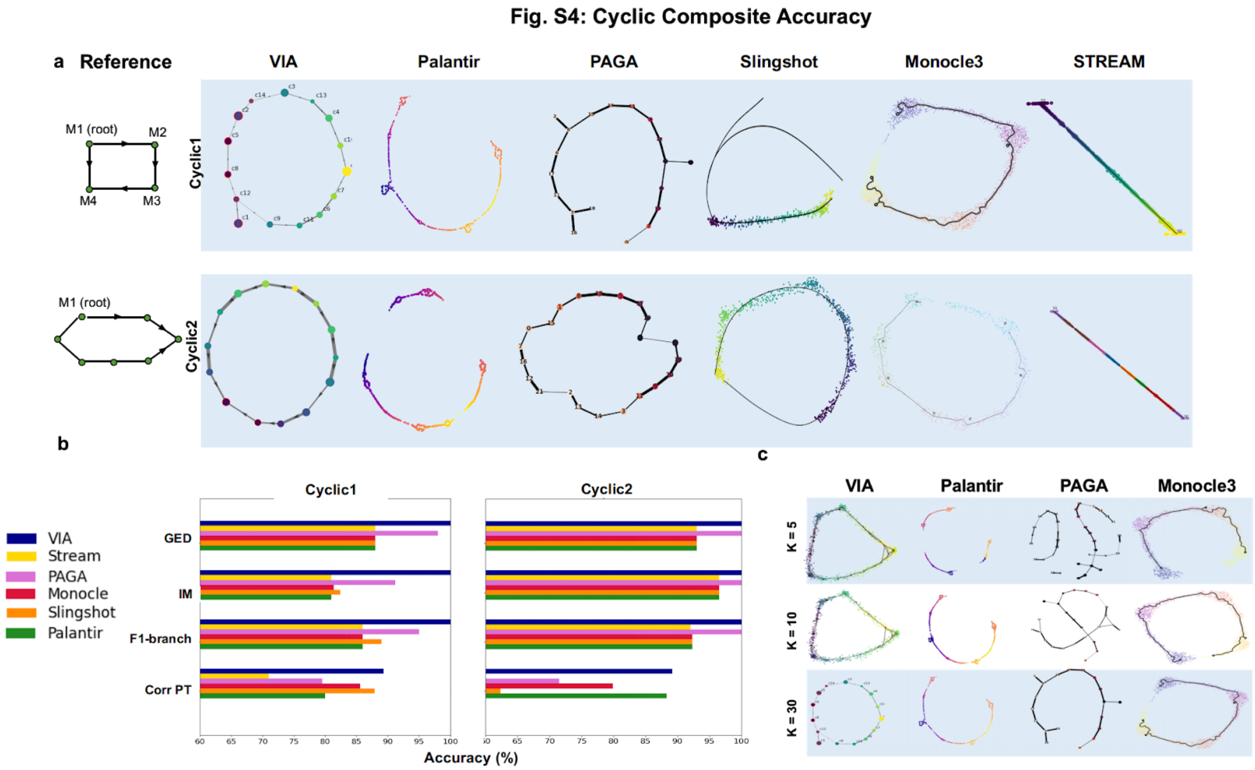


**Fig. S4 Benchmarking on cyclic synthetic datasets. (a)** Sample outputs by each TI method are shown for two cyclic datasets. The reference topologies are shown on the left. **(b)** The accuracy scores of 4 individual metrics (GED, IM, F1-branch and pseudotime correlation (Corr PT) comprising the composite accuracy score are shown individually. Note that there is no terminal cell fate due to the cyclic nature. **(c)** The cyclic dataset is more vulnerable to fragmentation for different levels of K in PAGA and Palantir, where at lower K (KNN), there is more fragmentation. This does not mean VIA is invariant to KNN, but that while the resolution (number of clusters present) in the graph changes, the overall topology remains intact. We notice that Monocle shows a linear topology (it does not close the loop) for either of the cyclic datasets (a). We show in **(c)** that increasing the number of KNN in Monocle3 does not necessarily close the loop either.

#### Fig. S5 Connected Composite Accuracy

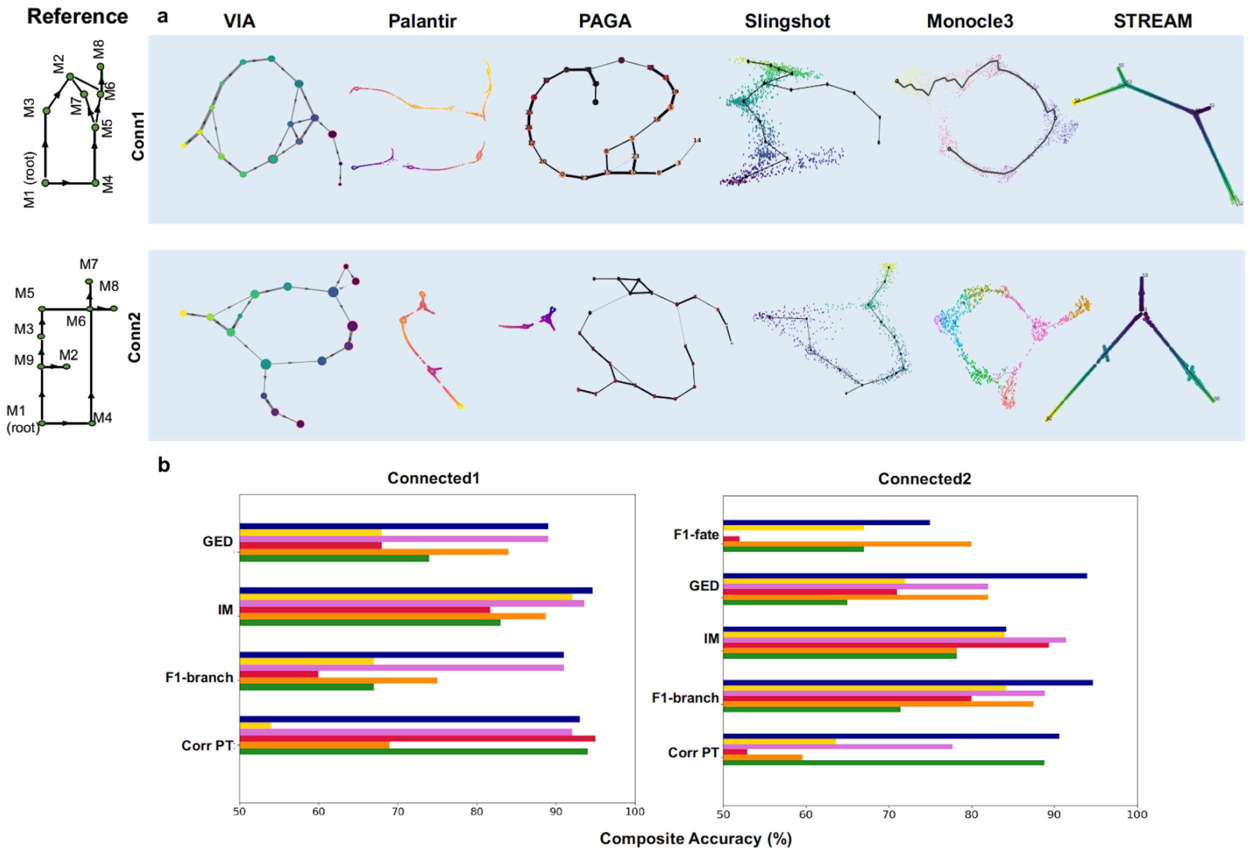


**Fig. S5 Benchmarking on connected synthetic datasets** **(a)** Sample outputs by each TI method shown for the connected datasets with reference topologies shown on the left. **(b)** The accuracy scores of each of the 5 individual metrics (F1-cell fate (F1-fate), GED, IM, F1-branch and pseudotime correlation (Corr PT)) comprising the composite accuracy score are shown individually.

### 9. VIA performance on real biological datasets

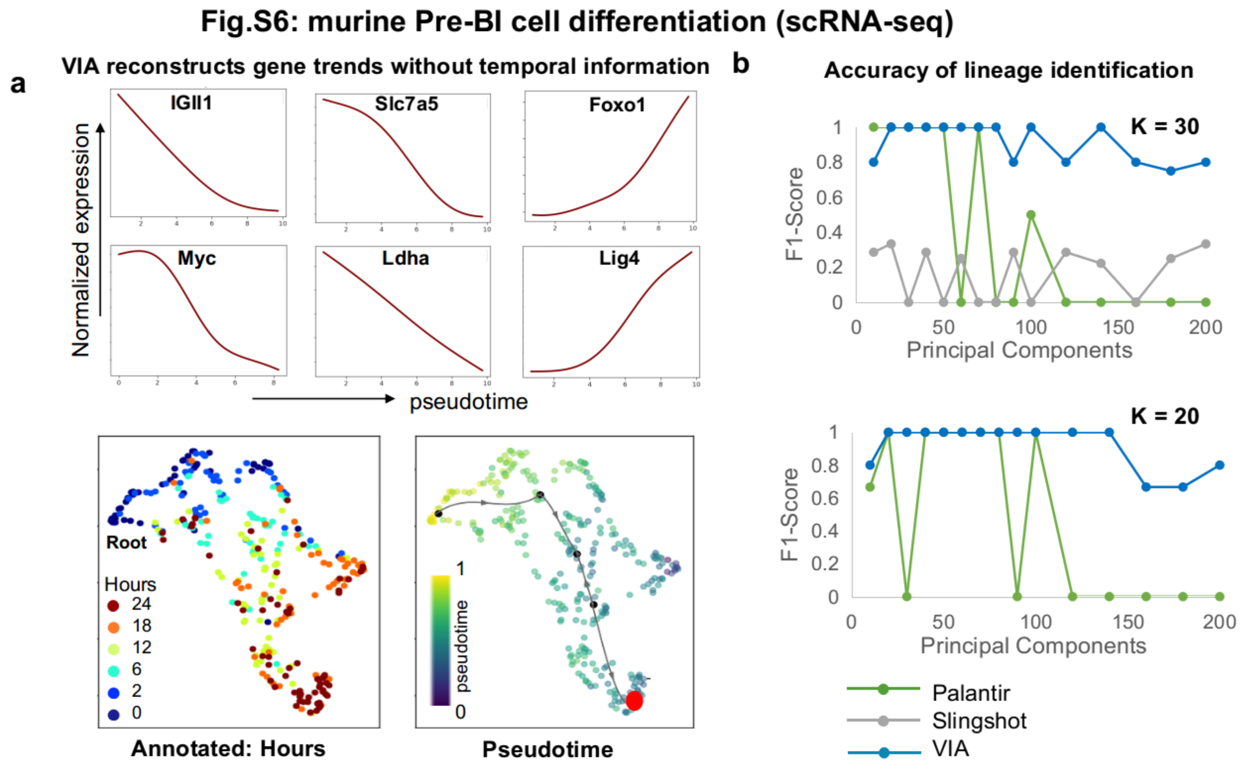


**Fig. S6 VIA analysis of mouse Pre-BI cell Differentiation[7] (scRNA-seq). (a)** VIA recapitulates up/down regulation of gene markers without prior knowledge of the temporal information. **(b)** Detection accuracy of cell fates (cells from 18-24 hours) across a range of PCs for K (nearest neighbors) = 30 (Top), and K = 20 (Bottom). Palantir performs comparably until the number of PCs exceeds ~50, after which it is unable to correctly identify later cell stages.

**Fig. S7: Human CD34+ Hematopoiesis (scRNA-seq)**

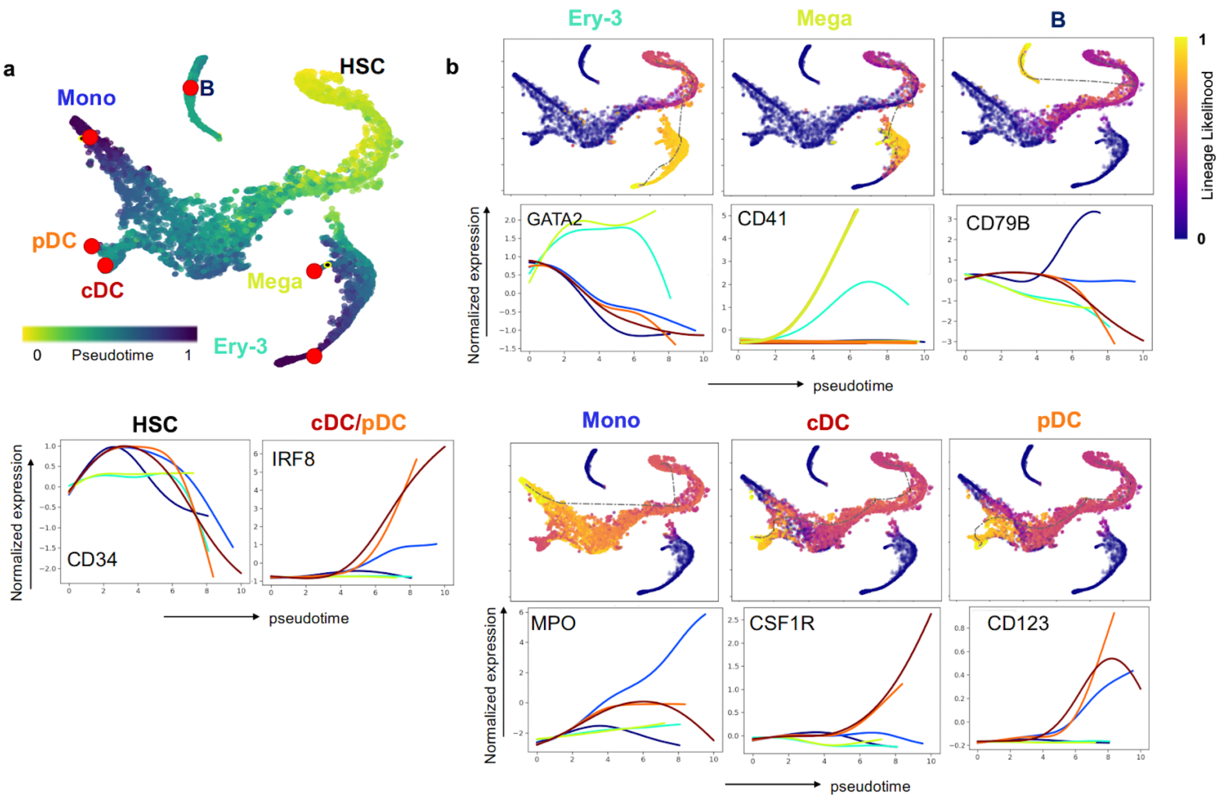


**Fig. S7 VIA analysis of human hematopoiesis[6] (scRNA-seq) (a)** We illustrate the detection of major (monocytes (Mono), erythrocytes (Ery) and B-cells (B)) and less populous lineages (megakaryocytes (Mega), conventional and plasmacytoid dendritic cells (cDCs, pDCs)). **(b)** (Top) UMAP embedding colored by the lineage likelihood (with the pathway overlaid) of the 6 identified terminal states. (Bottom) The gene expression trend of known marker genes along each lineage (including HSC (left)). Notably, the cDC and pDC populations show elevated *CSF1R* and *CD123* levels relative to other lineages. (see also the elevated level of *IRF8* in both cDC and pDC (left)), and the megakaryocyte population has higher *CD41*. The gene trend curves follow the same color code as the labels in (a).

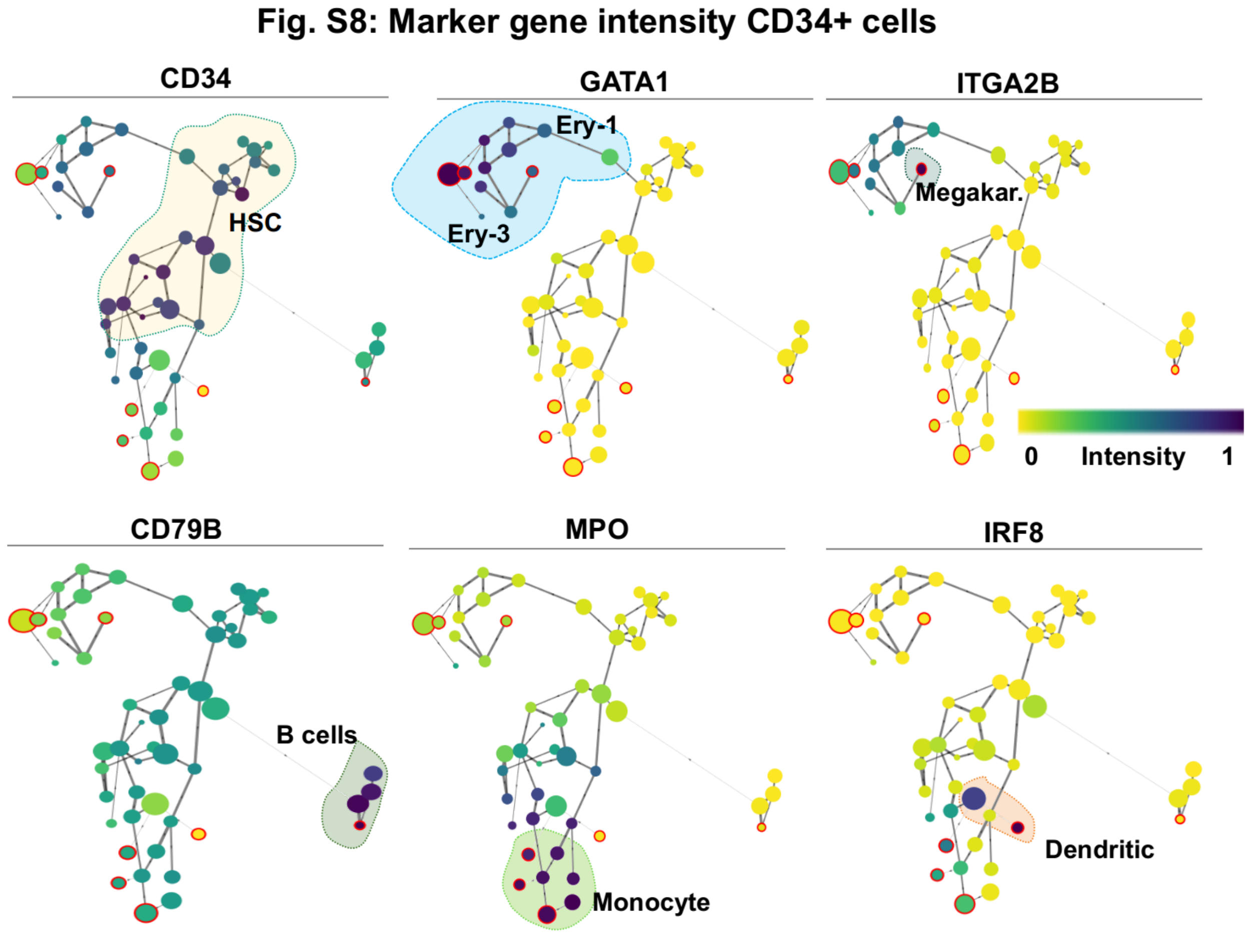


**Fig. S8: Lineage marker intensity along the VIA graph topology of human hematopoietic scRNA-seq cells.** Marker gene expression along the graph shows the location where gene expressions are up and down regulated for each of the major trajectories.

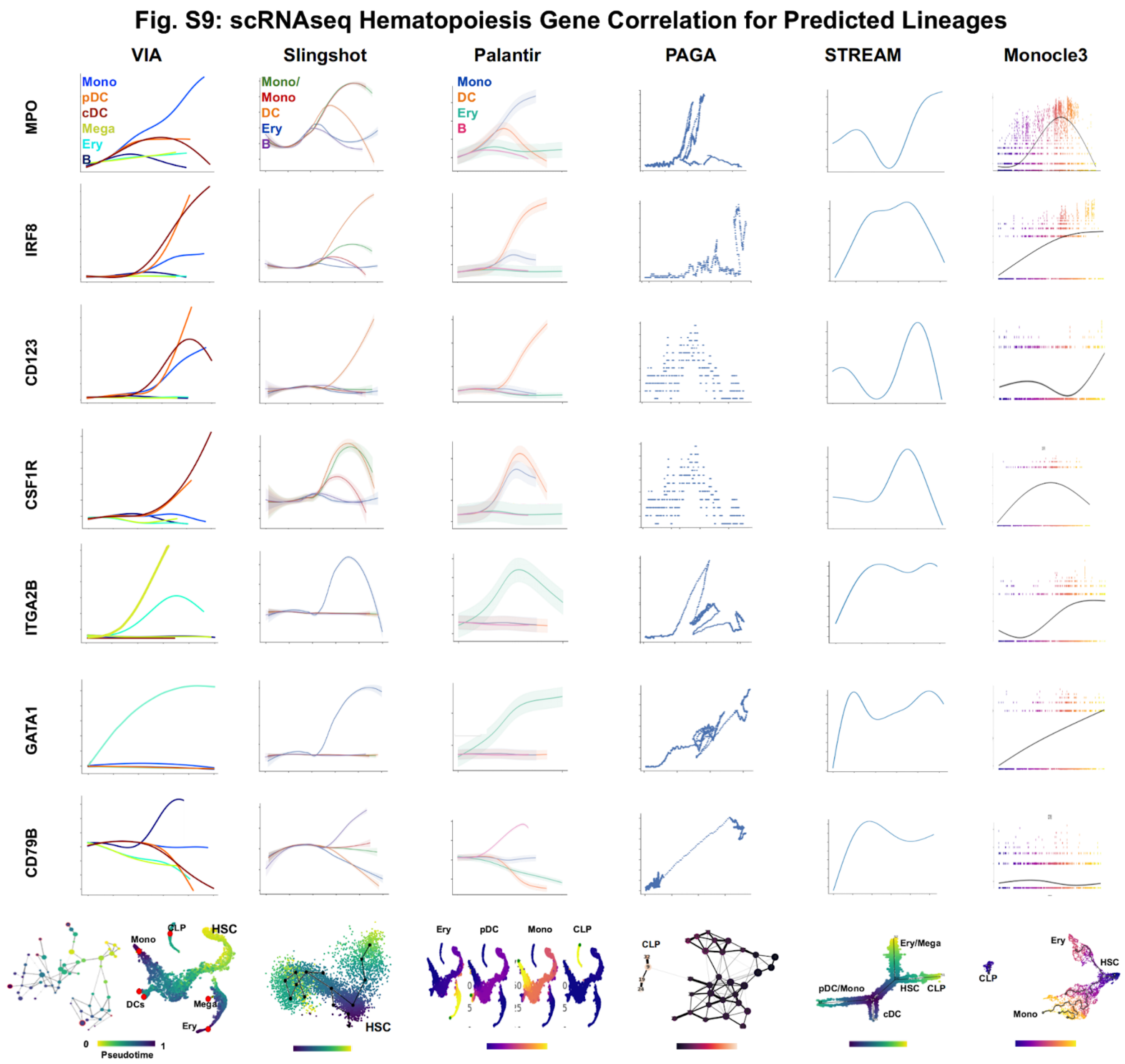


**Fig. S9 Comparison of marker gene correlation computed by different TI methods (for predicted lineages in scRNA-seq hematopoiesis).** Automated gene expression plots as a function of pseudotime for the lineages associated with 6 specialized cell types. For PAGA and STREAM we manually select the order of clusters/branches towards a terminal state. PAGA uses a sliding window approach to compute the expressions based on the selected clusters, whilst for STREAM we apply GAMs to plot the expression. For Slingshot's topology, the black lineage curves cannot be aligned with the t-SNE embedding, but we show the colored temporal ordering of cells on the t-SNE for ease of comparison to other methods.

#### Fig.S10: scATAC Hematopoiesis

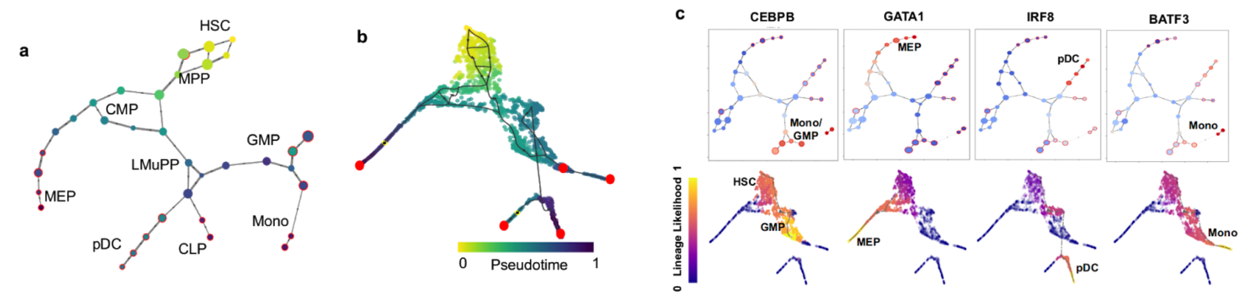


**Fig. S10 VIA on scATAC-seq human hematopoiesis**[13] **(a)** VIA graph and **(b)** UMAP colored by pseudotime using Buenrostro[13] pre-processing protocol. **(c)** Graph topology colored by marker TF motif accessibility locates the stage where changes in expression are triggered. (top) and lineage probability of pathways (bottom) reveal the correct order of cell type progression towards the differentiated state.

#### Fig. S11: scATACseq Hematopoiesis Gene Correlation for Predicted Lineages

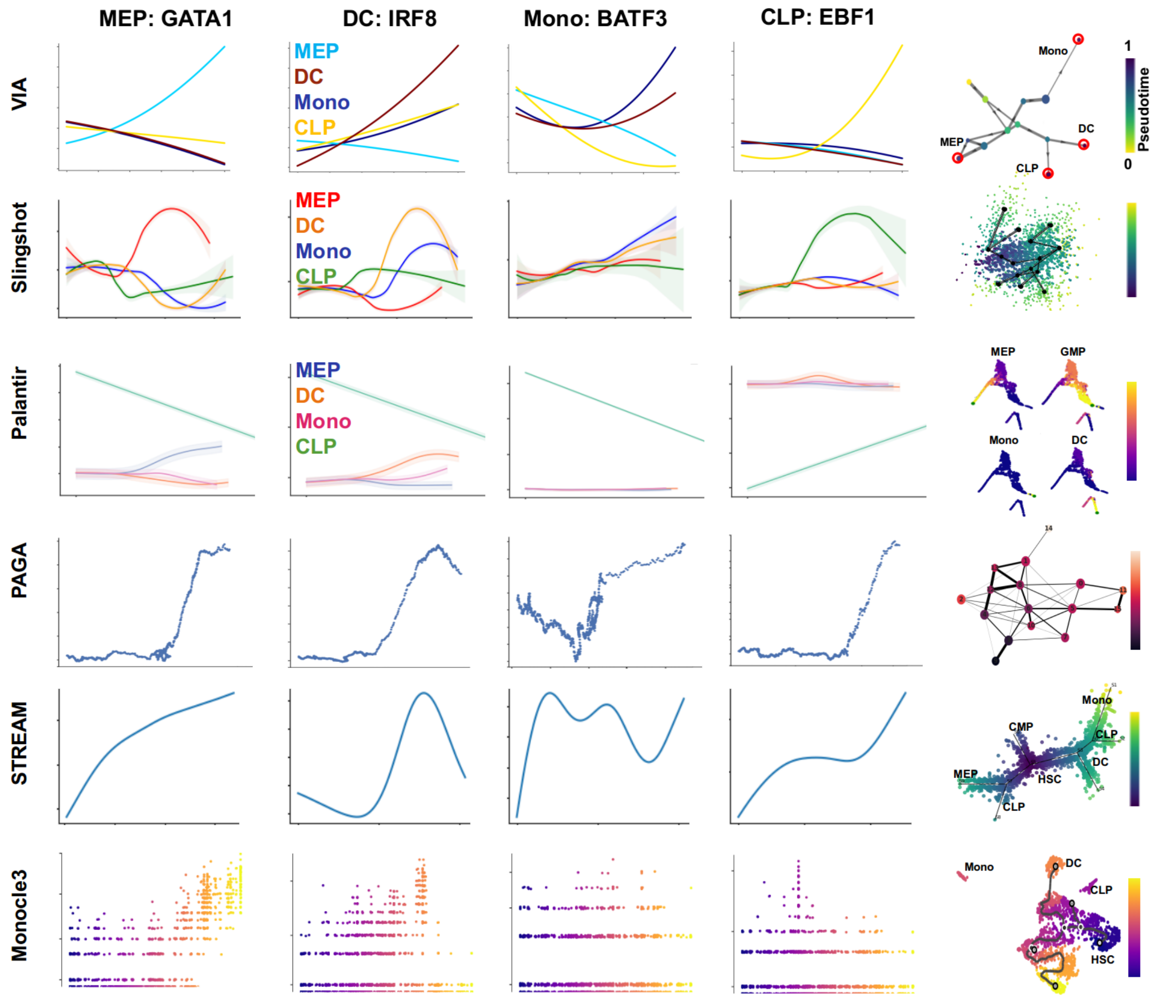


**Fig. S11 Comparison of marker gene correlation computed by different TI methods (for predicted lineages in scATAC-seq hematopoiesis).** Automated gene expression plots as a function of pseudotime for the lineages associated with CLP, MEP, Monocyte and DC committed cells. For PAGA and STREAM we manually select the order of clusters/branches towards a terminal state. PAGA uses a sliding window approach to compute the expressions based on the selected clusters, whilst for STREAM we apply GAMs to plot the expressions. Monocle3 did not plot fitting curves on this dataset.

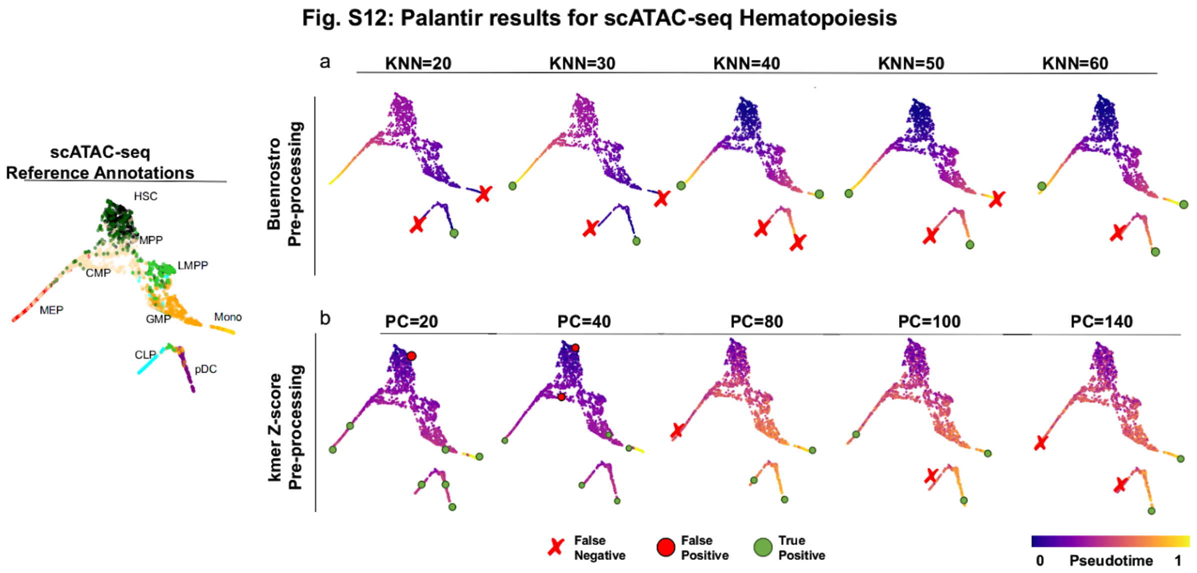


**Fig. S12 Palantir results for scATAC-seq hematopoiesis:** UMAP projection of single cells colored by annotated cell types **(a)** Palantir sample outputs for varying the number of K (nearest neighbors) when using principal components (PCs) from the original Buenrostro paper[13]. Start cell corresponds to an HSC cell. Palantir typically only finds the Erythroid lineage and one of the Monocytic or dendritic lineages **(b)** Palantir's accuracy on the kmer Z-score based PCs is better than in **(a)** but detection accuracy drops when the number of PCs increases.

#### Fig. S13: VIA on scRNA-seq Embryoid Body

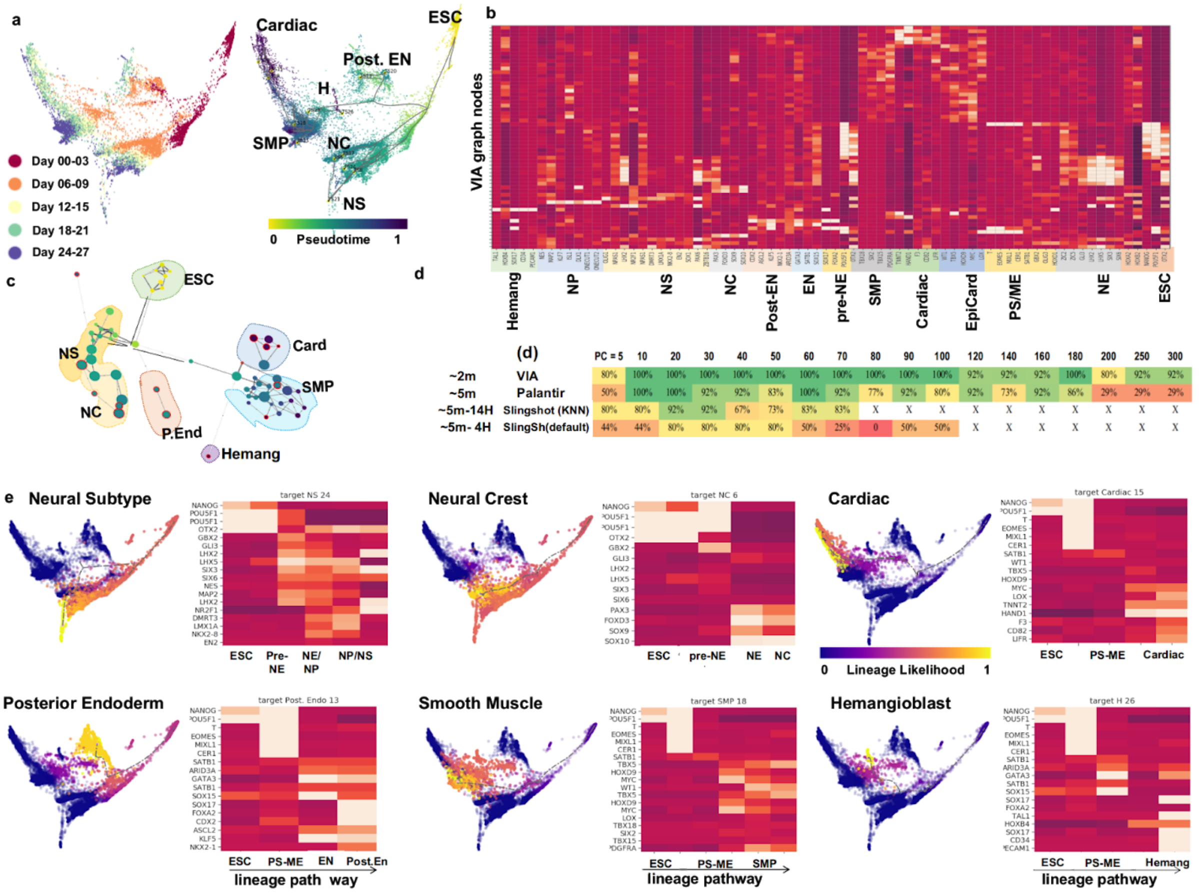


| | (d) | PC = 5 | 10 | 20 | 30 | 40 | 50 | 60 | 70 | 80 | 90 | 100 | 120 | 140 | 160 | 180 | 200 | 250 | 300 |
|---|---|---|---|---|---|---|---|---|---|---|---|---|---|---|---|---|---|---|---|
| ~2m | VIA | 80% | 100% | 100% | 100% | 100% | 100% | 100% | 100% | 100% | 100% | 100% | 92% | 92% | 92% | 100% | 80% | 92% | 92% |
| ~5m | Palantir | 50% | 100% | 100% | 92% | 92% | 83% | 100% | 92% | 77% | 92% | 80% | 92% | 73% | 92% | 86% | 29% | 29% | 29% |
| ~5m-14H | Slingshot (KNN) | 80% | 80% | 92% | 92% | 67% | 73% | 83% | 83% | X | X | X | X | X | X | X | X | X | X |
| ~5m- 4H | SlingSh(default) | 44% | 44% | 80% | 80% | 80% | 80% | 50% | 25% | 0 | 50% | 50% | X | X | X | X | X | X | X |

**Fig. S13: VIA analysis of human embryoid body[8] (scRNA-seq). (a)** (left) PHATE embedding colored by Day. (right) embedding colored by VIA pseudotime. VIA identifies 6 cell fates (arising from embryonic stem cells (ESCs)) labeled by corresponding cell type i.e. neuronal subtypes (NS), hemangioblast (Hemang), neural crest (NC), smooth muscle precursors (SMPs), cardiac (C) and posterior endoderm (EN), based on their gene expressions, concurring with the lineages identified by Moon et al.[8] Other abbreviations of intermediate states include: neuroectoderm (NE), neural progenitors (NPs). epicardial precursors (EpiCard), primitive streak (PS), mesoderm (ME). **(b)** Heatmap of marker gene expressions to identify cluster cell-type. **(c)** Cluster-level topology of ESCs differentiating to the main lineages. **(d)** F1-Score of lineage prediction accuracy corresponding to the cell fates for Palantir, Slingshot and VIA across a wide range of input Principal Components (PCs). Palantir and Slingshot do no not capture the cardiac cell fate; Slingshot also misses the neural crest. Runtime of each method is highlighted on the left side of the chart. Slingshot is not feasible for higher numbers of PCs as the runtime becomes prohibitive (>14 hours) **(e)** PHATE embedding colored by VIA's lineage likelihood shows the differentiation pathway towards specialized cells. VIA also produces a cluster-level pathway of lineage relevant clusters that can be used to generate lineage heatmaps. For instance, the neural subtype traces a path from the ESC cells, followed by pre-neural ectoderm (pre-NE), neural ectoderm and neural progenitors, ending at a neural subtype.

#### Fig. S14: VIA on scRNA-seq endocrine genesis

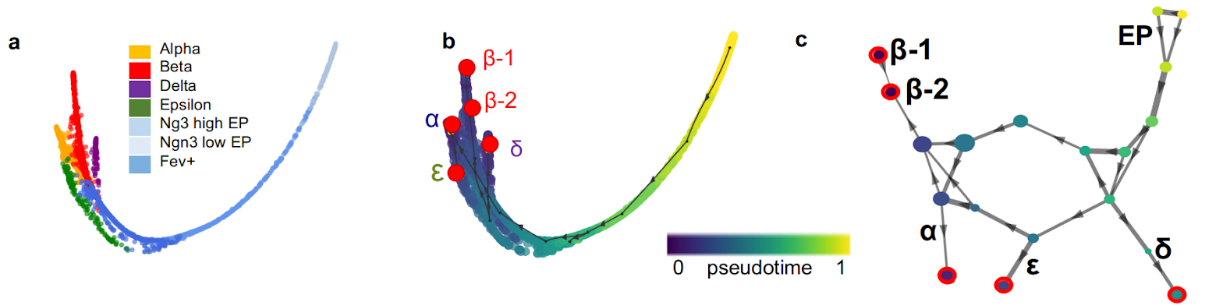


**Fig. S14 VIA analysis of endocrine differentiation[9] (scRNA-seq) (a)** PHATE embedding colored by known labels. **(b)** VIA-inferred pseudotime projected on the PHATE embedding. Five terminal states, i.e. Alpha, Beta-1, Beta-2, Delta and Epsilon lineages with Beta-2 being an *Ins2+* Beta subtype, are highlighted. **(c)** VIA graph colored by pseudotime and terminal states marked in red outlines.

**Fig. S15: Marker Gene Correlation for Predicted Pancreatic Endocrine Lineages**

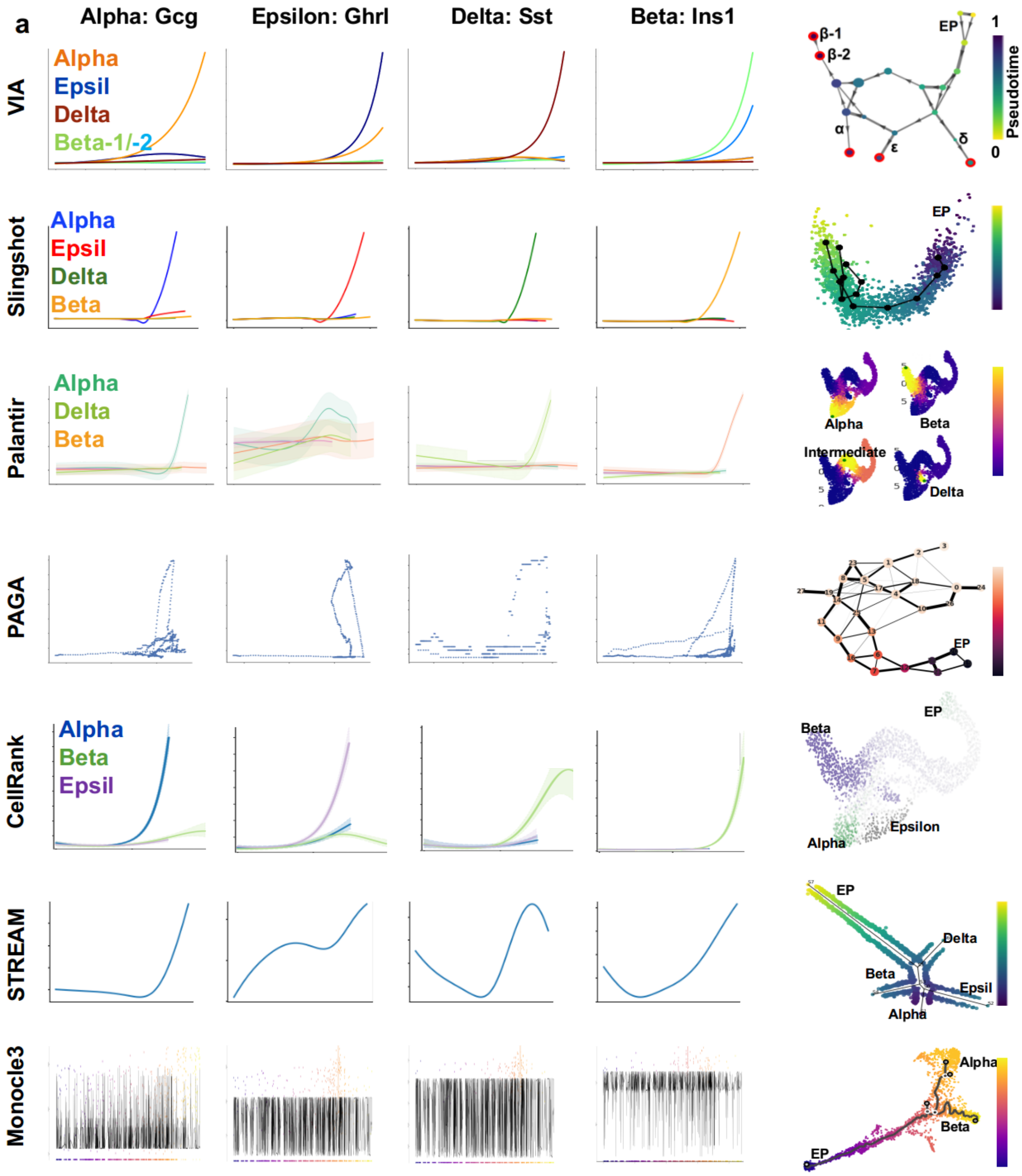


**Fig. S15 Comparison of marker gene correlation computed by different TI methods (for predicted endocrine lineages).** Automated gene expression plots as a function of pseudotime for each of the 4 pancreatic islets: Alpha (Gcg), Beta (Ins1 and Ins2), Delta (Sst) and Epsilon (Ghrl). For PAGA and STREAM we manually select the order of clusters/branches towards a terminal state. PAGA uses a sliding window approach to compute the expressions based on the selected clusters, whilst for STREAM we apply GAMs to plot the expressions. The dark curves on the Monocle plots are very noisy and overwhelm the original data points.

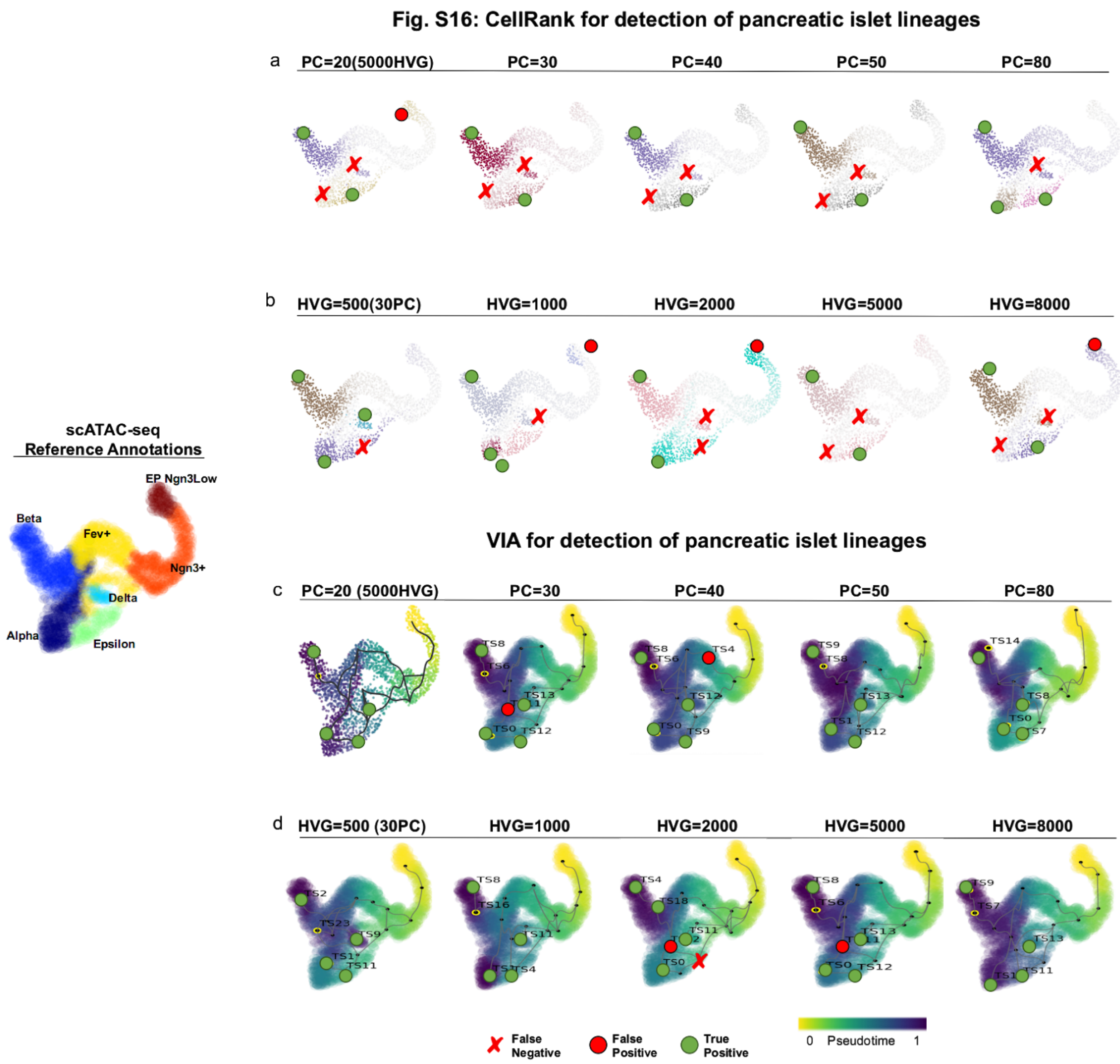


**Figure S16 CellRank for endocrine genesis (scRNA-seq):** UMAP projected cells colored by annotated cell type **(a)** CellRank sample outputs when the number of PCs is changed shows that the Alpha and Epsilon islets tend to be merged and the Beta and Delta similarly merged. **(b)** This holds for when the number of highly variable genes (HVG) is changed. **(c)** VIA sample outputs when the number of PCs is changed shows that the 4 islets are consistently detected. The Ins2+Ins1- Beta-2 population (small yellow dot) is also detected. **(d)** This holds for when the number of highly variable genes (HVG) is changed.

### Supplementary Note 4: Noise and drop-out in scRNA-seq data

We use the R package "seqgendiff" (Gerard, 2020) to add noise to the real RNA-seq datasets to intensify the attributes of noise in real data. Seqgendiff uses binomial thinning to subsample counts using the binomial distribution and reduce the total count expression by a specified factor. Before applying binomial thinning, seqgendiff randomly selects a subset of the original features (genes) so that the results are not dependent on the behaviour of a few features (genes).

We first show the deterioration in VIA's inferred topology for a synthetic 4-leaf multifurcation **(Fig. S17)**, when the level of noise is increased (or the signal strength is diminished) by a factor of 2 at each step. Note that prior to binomial thinning, only 10% of the original features are retained (at random) from the original input to mitigate dependency on a few select features. Since the signal in the synthetic data is strong, we can recover the overall topology and delineate 3-4 of the cell fates until a noise-factor of around 5. At higher levels of signal atrophy (Factor 7 and 8), the true structure is imperceptible.

To show the impact of adding noise to a real dataset (such as the endocrine dataset where the Delta lineage is very small, and the Epsilon lineage is easily merged with the Alpha islet cells), we compare VIA to other methods in terms of the inferred topology and predicted gene trends which are computed based on the single-cell level lineage probabilities and cell ordering. The UMAP embeddings in **Fig. S18b-c** illustrate the level of added noise showing the separation between lineages becoming obscured in the noised-data visualization. A thinning factor = 1 was used on 25% of genes as adopted in Gerard 2020 and the package vignettes. VIA's inferred trajectory remains true to the expected progression from endocrine progenitors, then to a forking at the Fev+ intermediate state towards different islets. More importantly, STREAM, Slingshot and Palantir appear to suffer most from the added noise as seen by the high 'cross-talk' between the plotted lineage trends (e.g. in the Slingshot trends), or the detection of only two islets (e.g. Palantir), or in the case of STREAM the compromised topology where Alpha/Beta cells are plotted along the same branch, and the Epsilon/Delta are likewise combined.

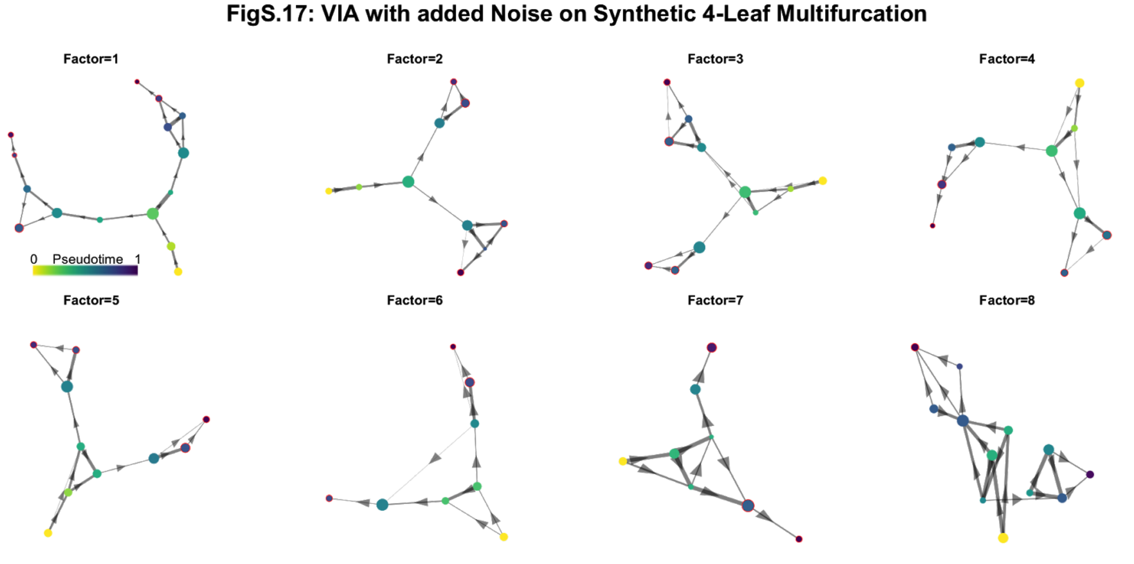


**Fig. S17 Impact of noise on VIA's inferred topology for a synthetic 4-leaf multifurcation.** Noise is added using R package "seqgendiff". 10% of the original 1000 features are sampled and retained. Then binomial thinning is applied where the aggregate signal is lowered to $2^{-Factor}$ of its original strength.

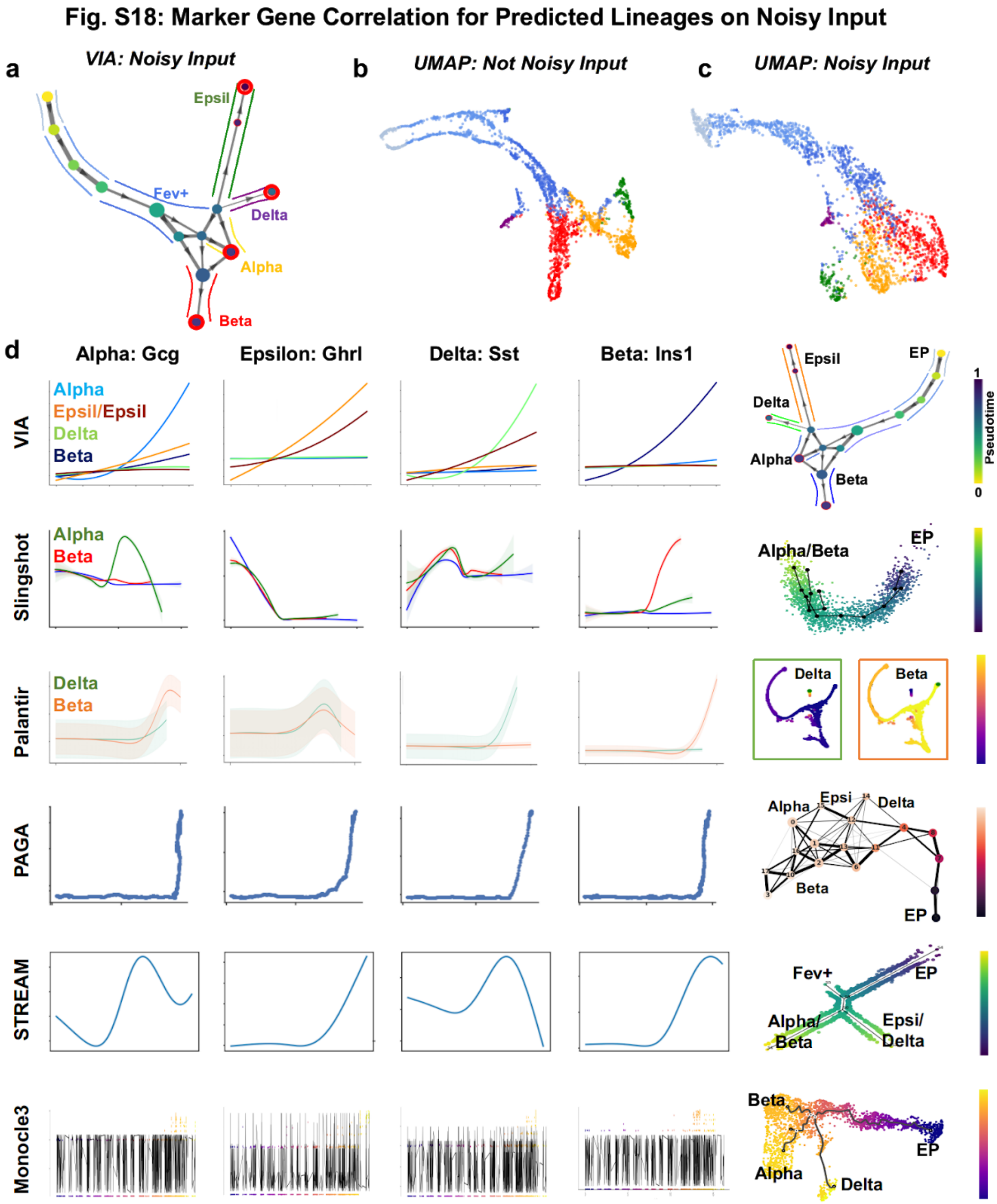


**Fig. S18 Gene correlation for predicted lineages on the "noised" pancreatic endocrine dataset.** 30% of quality filtered genes are retained, followed by factor 1 binomial thinning **(a)** VIA topology for noised input shows the desired progression of cells through 2 channels of Fev+ cells towards the 4 islets. **(b)** UMAP of original input **(c)** UMAP after applying binomial thinning blurs boundaries between the islets **(d)** comparison of inferred topology and gene trends shows that many methods merge lineages together and fail to detect 4 distinct islets corresponding to the Alpha, Beta, Delta and Epsilon lineages.

#### Fig. S19 Cardiac progenitor (scRNA + scATAC)

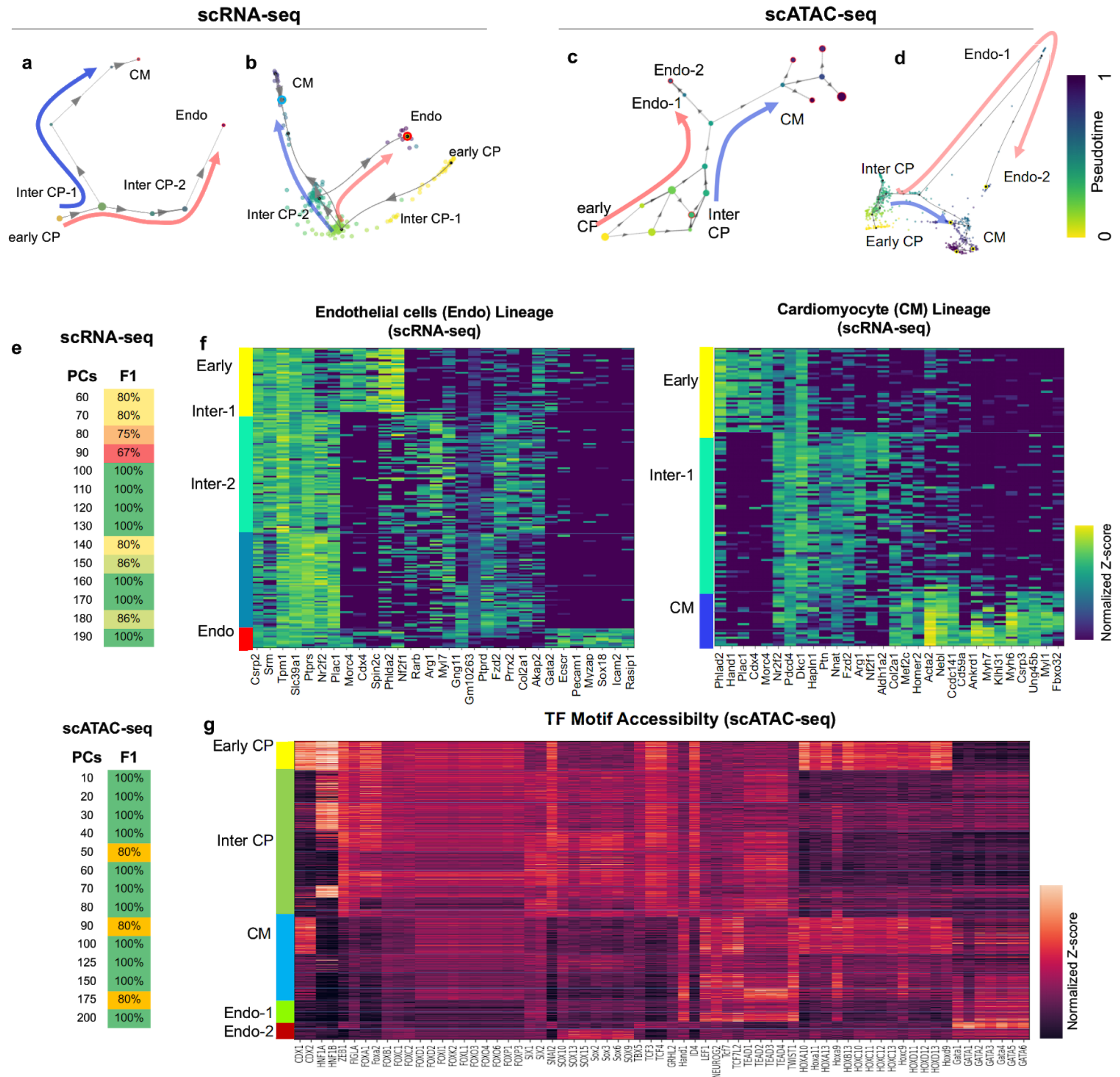


**Fig. S19 VIA analysis of *Isl1*+ Cardiac progenitor scATAC-seq and scRNA-seq**[14] **(a)** VIA graph colored by pseudotime for scRNA-seq data. Bifurcation of the pluripotent *Isl1+* CPs towards endothelial (Endo) (red path) and cardiomyocyte (CM) fates (blue path). Cell type labels correspond to annotations by Jia et al.[14]. **(b)** Single-cell UMAP embedding of scRNA-seq data. The graph level lineage paths (grey lines) and pseudotime are projected on the embedding. **(c)** VIA graph based on scATAC-seq, colored by pseudotime. Nodes labeled by majority cell type based on annotations. Again a bifurcation after the intermediate CPs shows a path towards CM (blue) and Endo (red) lineages. **(e)** F1-accuracy for detection of the CM and Endo lineages for variable number of PCs. The low cell count of the scRNA-seq dataset (197 cells) may account for slightly lower accuracy compared to the larger scATAC-seq dataset of 695 cells. **(f)** Top 5 most differentially expressed genes for each VIA cluster-node for each lineage in the scRNA-seq dataset (sidebar colored by majority annotation, following the color code in **Fig. 2d**). **(g)** Cluster level TF motif accessibility across TF motifs shortlisted by Jia et al.

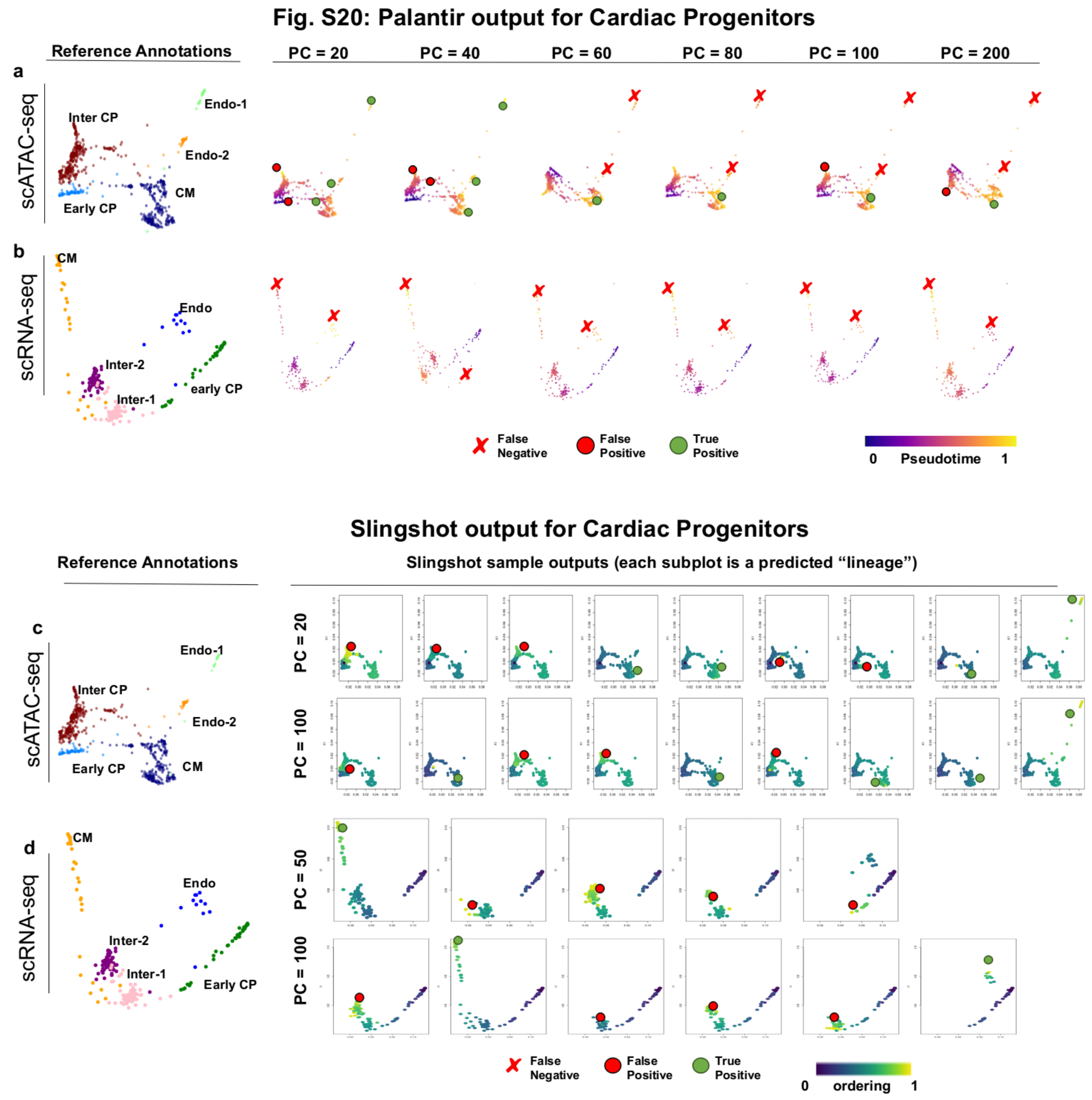


**Fig. S20 Palantir and Slingshot on Cardiac Progenitors (a)** scATAC-seq (695 cells annotated by cell type) of Cardiac Progenitor (CP) lineage commitment. Start cell chosen as Early CP. Palantir sample outputs colored by pseudotime for when varying the number of Principal Components (PCs) At PC=60 and above, Palantir misses the Endothelial cell fates **(b)** scRNA-seq (197 cells annotate by cell type) for different number of PCs misses the bifurcation to endothelial and cardiomyocyte cell fates. **(c)** Sample outputs for Slingshot (root cell selected as cluster corresponding to Early CPs) shown for 20 and 100 Principal Components (PCs). Each subplot is a detected lineage. At PC=20, 9 lineages are detected (colored by lineage likelihood) of which 5 are false positives (corresponding to intermediate CPs) **(d)** scRNA-seq (197 cells annotate by cell type) of CP differentiation shown for PC=50, 100 for Slingshot. Early CP set as the start state. 2-3 of the 5 detected lineages are false positive, corresponding to intermediate CPs

#### Fig. S21: STREAM output for Cardiac Progenitors

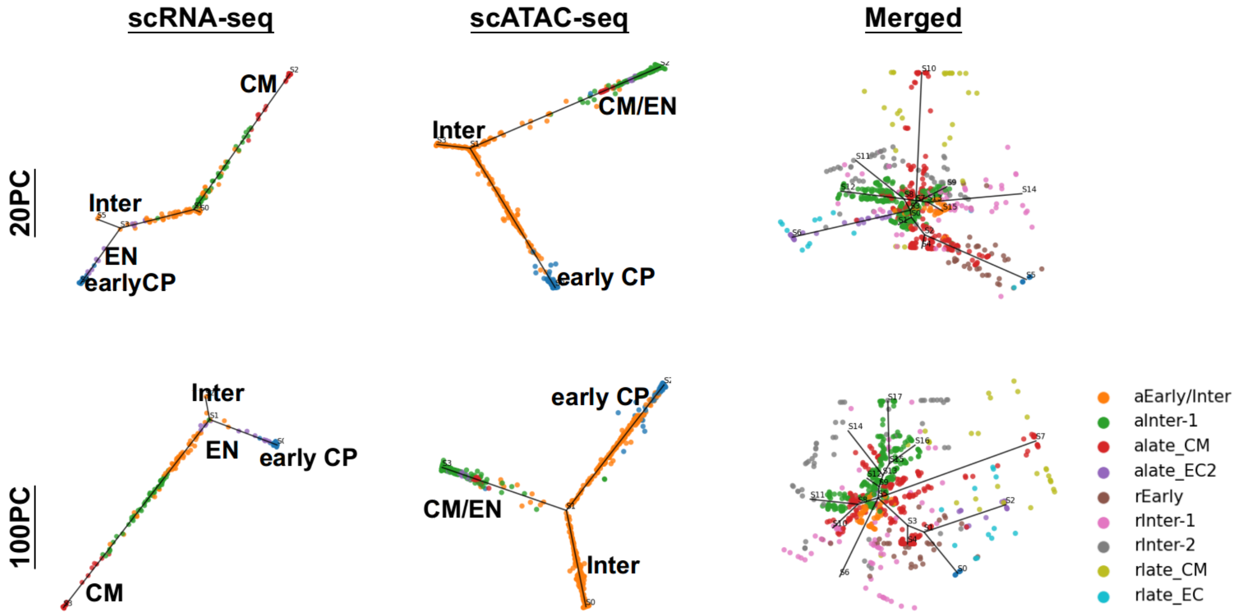


**Fig. S21 STREAM on Cardiac Progenitors:** The sample outputs of STREAM's topology colored by reference cell type at 20 and 100 PCs for each of the three types of data (scRNA-seq, scATAC-seq and merged), illustrates that the bifurcating structure is not correctly captured. In the scRNA-seq, we see that the small branch shortly after Early Cardiac Progenitors (CPs) is actually an intermediate population. The Endothelial cells are placed just after the CPs. In the scATAC-seq, the first branch after the early CPs is again populated by intermediate cells, and the cardiomyocytes (CM) and endothelial (EN) cells are merged together on the subsequent branch. The merged data is quite difficult to interpret as almost each cell type is assigned a unique branch.

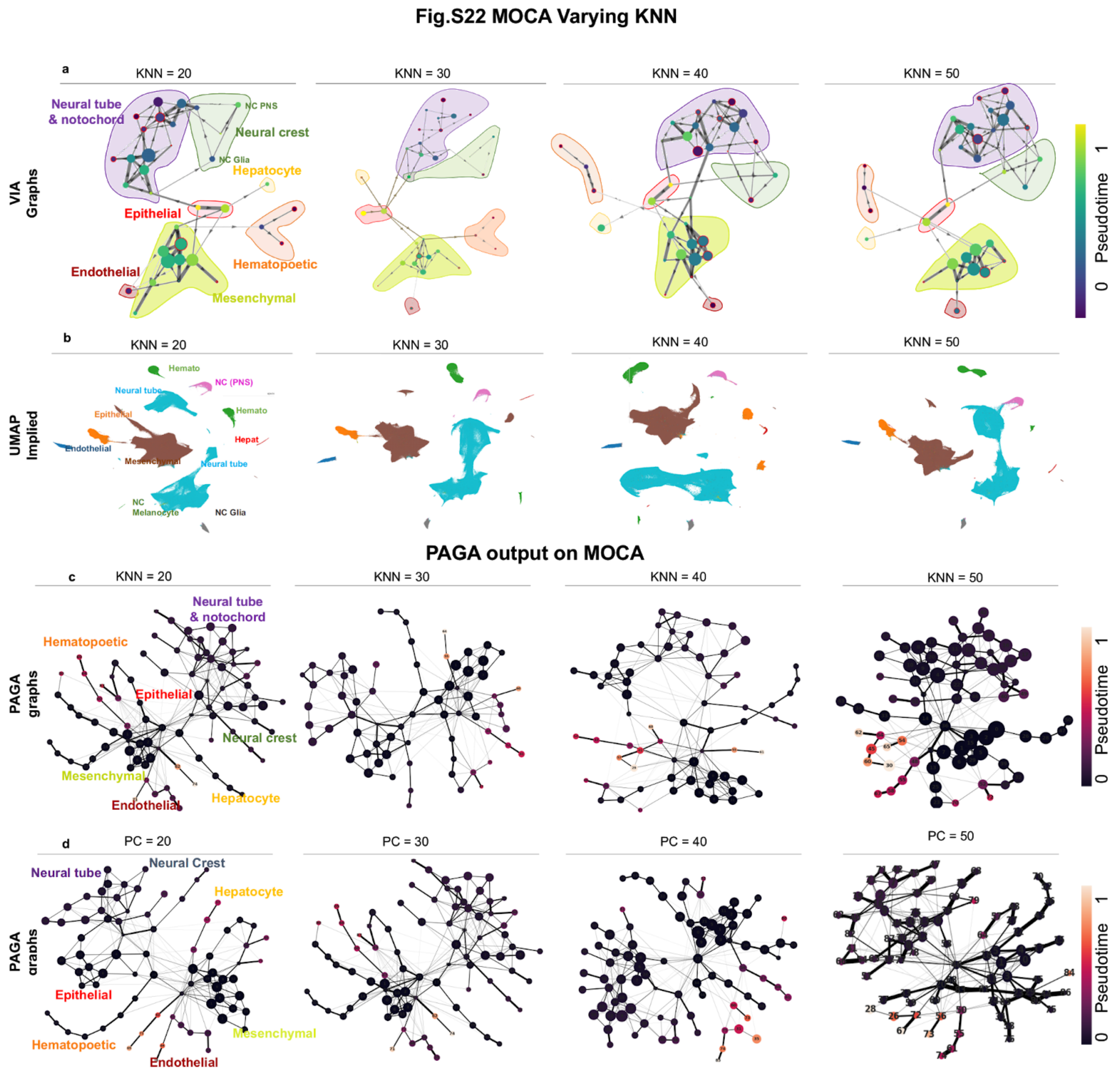


**Fig. S22 MOCA sample outputs across benchmarked methods (a)** VIA graph colored by pseudotime of 1.3 Million cells MOCA when the number of K (nearest neighbors) is varied. Key neighborhood relationships are maintained and pseudotime exhibits an salient gradient at high cell counts **(b)** UMAP implied (disconnected) projections that would be input to Monocle3 or Slingshot for TI show that the major trajectories and their relative placement are subject to fragmentation and random placement when the K varies. **(c)** PAGA graphs colored by DPT pseudotime show a highly connected structure that remains stable across parameter configuration. Pseudotime gradient is distorted.

#### Fig. S23: FACED - other TI method outputs

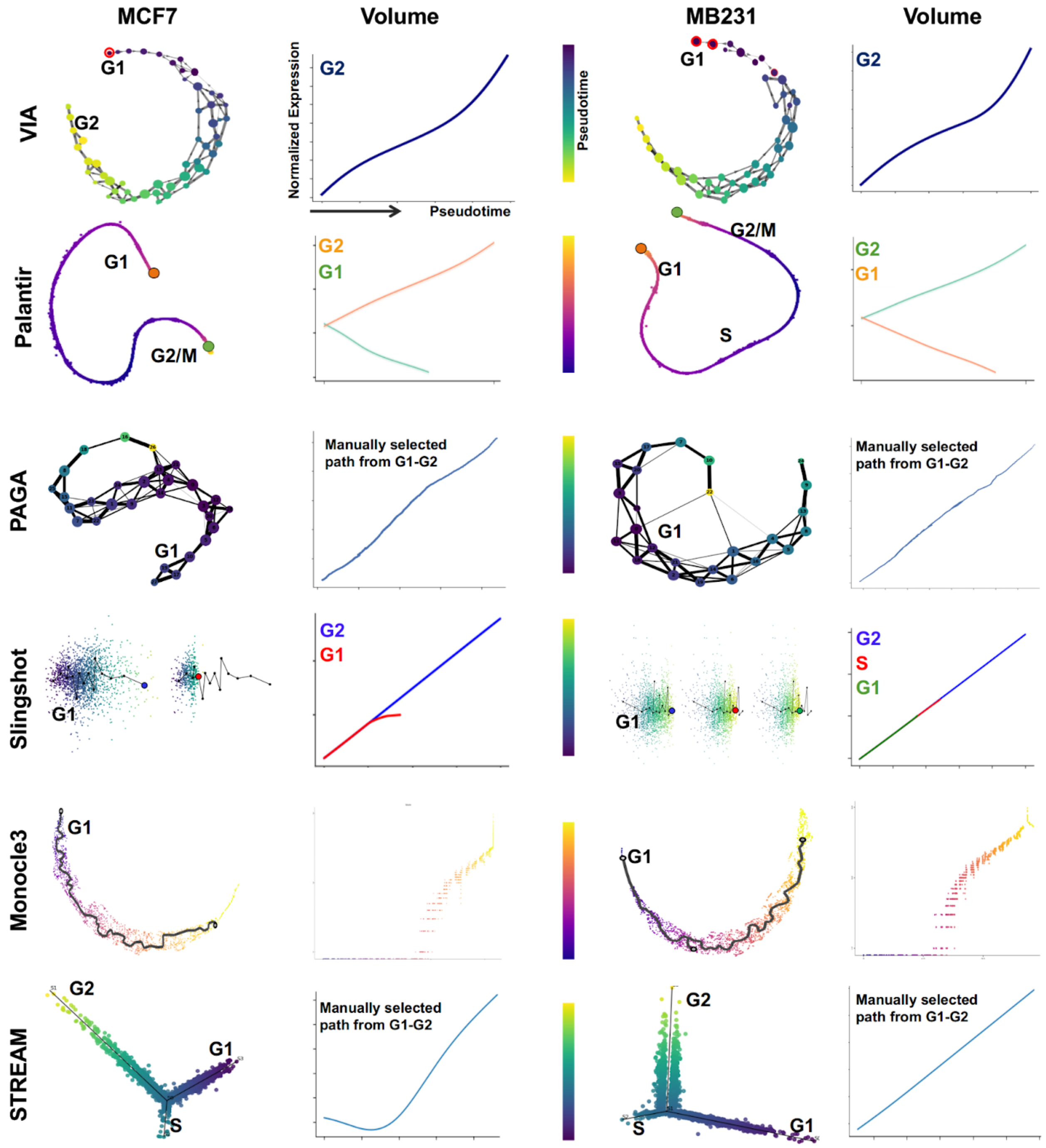


**Fig. S23 FACED biophysical profiles of cell-cycles analyzed by other methods.** The topologies and the temporal trend of volume inferred by other methods is shown for MCF7 and MDA-MB231. Note that for PAGA and STREAM we manually select the branches or clusters used in the trend plots. In STREAM the S cells are placed in a separate branch in a bifurcating structure.PAGA structure has an "S-shape" which we do not follow when selecting suitable clusters from G1 to G2 state. Many methods such as Slingshot and Palantir show multiple lineages ending at different stages along the cell cycle even though the root cell is selected as a G1 stage at the 'tail end' of the data representation.

#### Fig. S24: Predicted and Actual DNA content

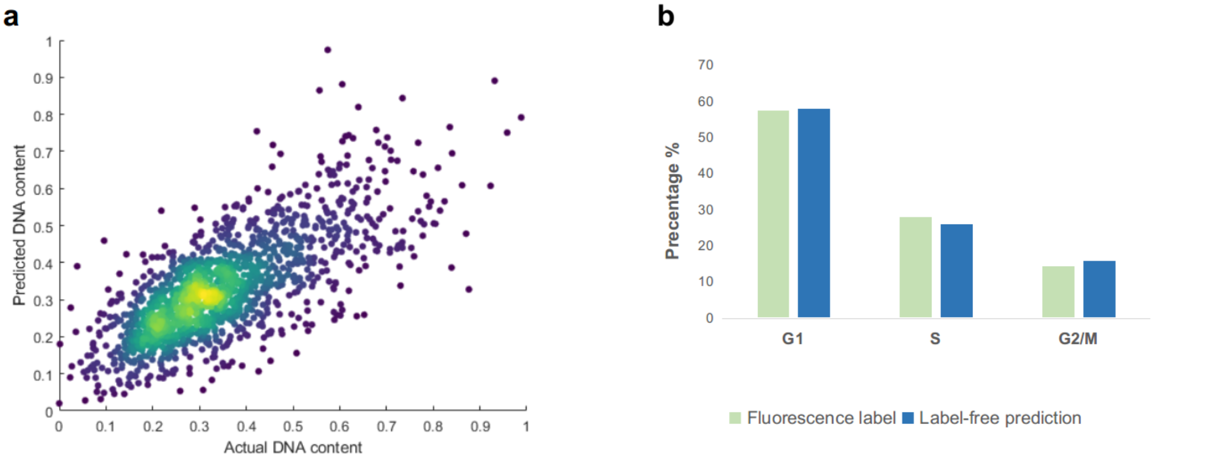


**Fig. S24 Validation of cell cycle analysis of breast cancer cells (including both MDA-MB231 and MCF7)**: **(a)** Correlation between the actual and predicted DNA content (by biophysical phenotypes extracted from FACED single-cell images. **(b)** Fractions of cells in G1, S and G2/M phases based on fluorescence label and label-free prediction, respectively.

#### Fig. S25: MB231 cell cycle TI w/o size related features

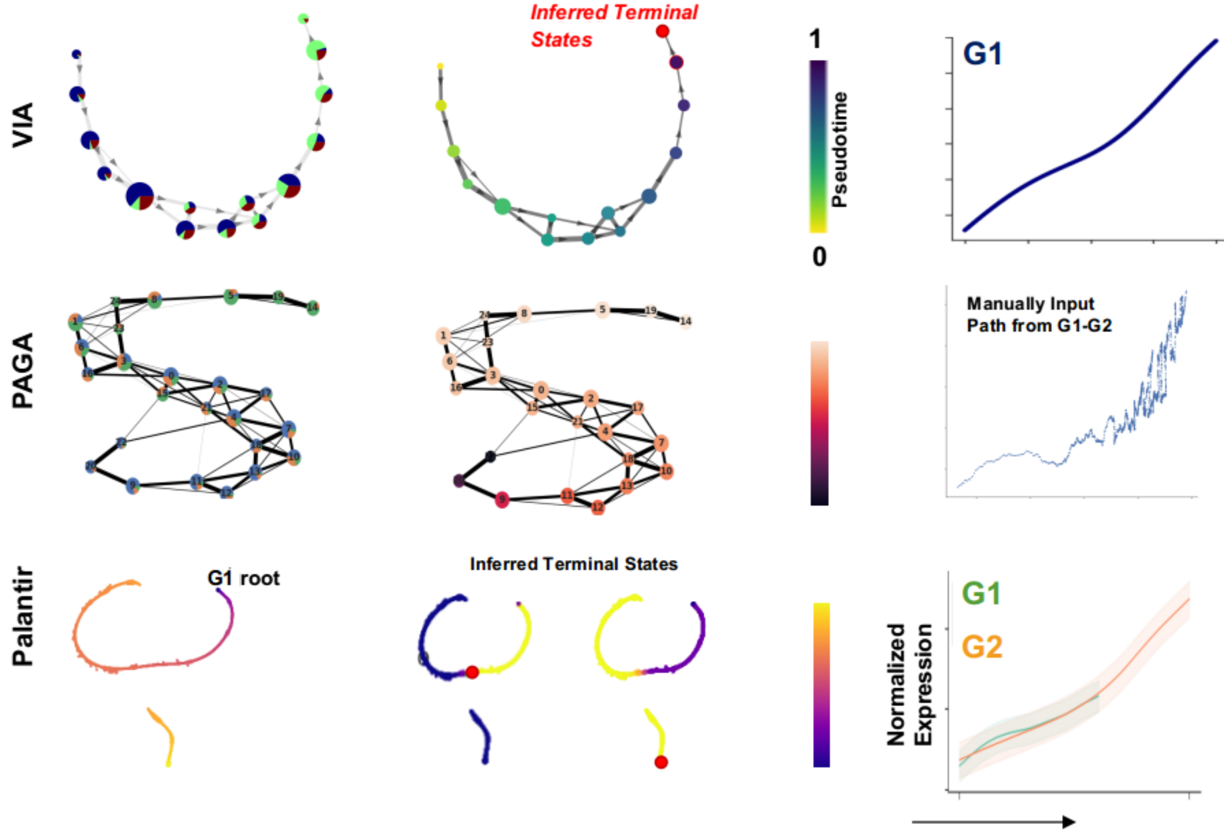


**Fig. S25 Cell-cycle TI predicted without cell volume (and volume-correlated) features (MDA-MB231 cells).** Volume and features $|Corr(feature, volume)| > 0.8$ are removed from the input to TI. We then plot volume to see if the correct trend along the inferred cell ordering is observed, including a slow-down in S-phase. Note that for PAGA, the clusters from a root cell to a final "G2 cluster" are manually provided.

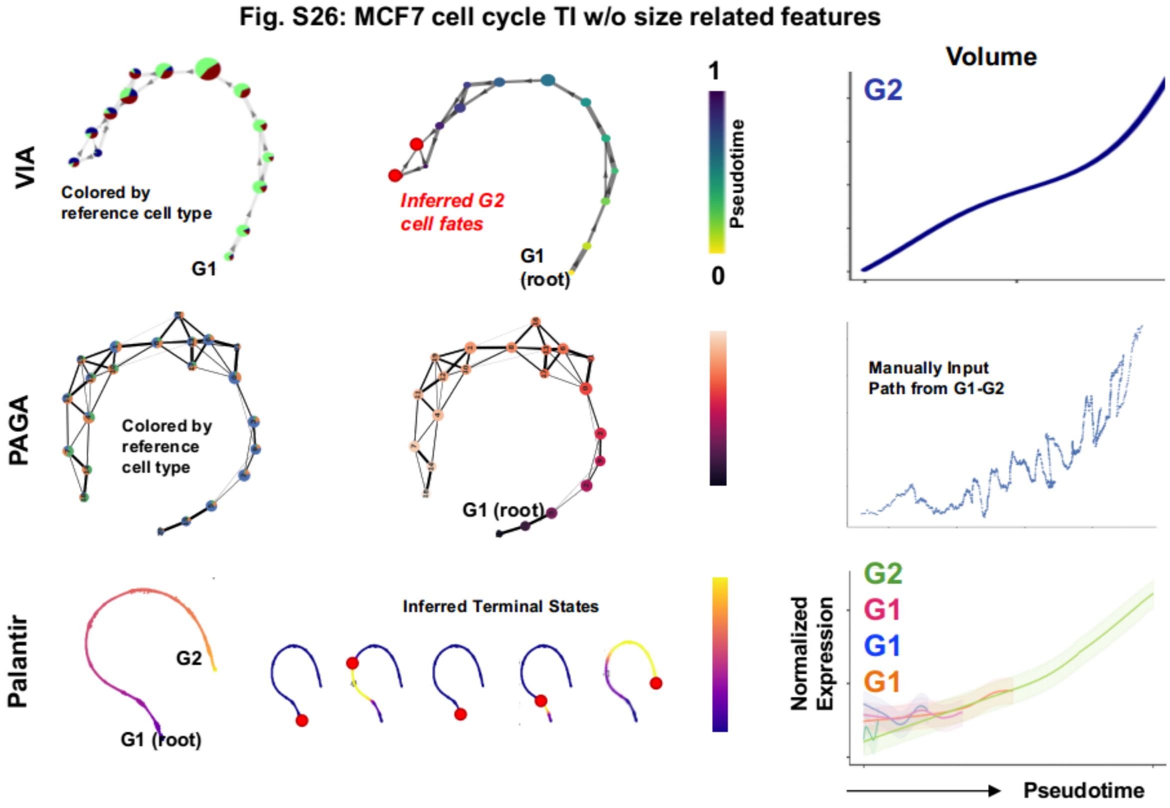


**Fig. S26 Cell-cycle TI predicted without cell volume (and volume-correlated) features (MCF7 cells).** Volume and features with an absolute correlation with volume above 80%, i.e. $|Corr(feature, volume)| > 0.8$ , are removed from the input to TI. We then plot volume to see if the correct trend along the inferred cell ordering is observed, including a slow-down in S-phase. Note that for PAGA, the clusters from a root cell to a final "G2 cluster" are manually provided.

### 10. Supplementary Note 5: VIA on gene matrix without dimensionality reduction

VIA handles a broad range of input feature dimensions that typically characterize various omic datasets. Imaging and mass cytometry features tend to remain within 10-100 features (antibodies or morphological features), whereas scRNA-seq produces 5000-15000 genes even for small sample sizes of hundreds or a few thousand cells. Due to the sparsity of count matrices, it is standard practice to filter the cells and genes according to certain thresholding criteria. However, even after such a step, the number of features can still be multiple magnitudes of the cell count and thus dimensionality reduction such as PCA is used to enhance the signal, before running further unsupervised computational analysis.

Several TI methods, including top-performing ones like Slingshot, STREAM and Monocle3 all require or highly recommend further reducing the dimensionality using a second method such as UMAP, TSNE or the like. As such, the ability of VIA to perform accurate TI on either a full feature set of 'unprocessed' cytometry features, or easily handle 100-200 PCs in the ATACseq and RNAseq datasets allows users to minimize the level of dimensionality reduction compared to other methods.

For instance, STREAM uses either Spectral Embedding, UMAP or MLLE after PCA which adds an additional layer of method choice and subsetting of components. While it is possible to override the second dimensionality step in STREAM, requiring it to skip the PCA is not practical and, taking 4.5 hours for a small dataset of 500 genes and 2000 cells. Monocle3 requires UMAP on a selected number of Principal Components, and subsequently retains the first 2 (default) or 3 components of the manifold embedding for TI. In Slingshot, the guidelines recommend only retaining the first 2-10 PCs. This ensures synchrony between the visualization of lineages and principal curves using the input data. In Slingshot, skipping the PCA step even for lower dimensional cytometry data (10-100 features) is not a viable option as the lineage curves cannot be well visualized on the raw input features and the runtime scales very poorly along the feature space. Other methods like Palantir which can receive raw features or a higher number of PCs involve steps within their algorithms that quite aggressively subset the number of diffusion components (default of 10 eigenvectors is computed and even fewer DCs) retained for further TI analysis.

We stretch the concept of VIA digesting even higher dimensional data by showing the VIA sensibly processes both real and synthetic data using several thousands of genes without any form of dimensionality reduction. This shows that in principle VIA can handle extremely high dimensions without distorting the data, a promising sign as the feasible dimensionality of flow and mass cytometry experiments grows, and the cell counts of transcriptomic experiments increases making it reasonable to retain more genes in downstream computational analysis.

We used the synthetic datasets spanning various topologies to observe how faithfully VIA uncovers topology, pseudotime and cell fate prediction. Each synthetic dataset comprises 1000-3000 'cells' across 1000 features. These features were provided to VIA without any dimensionality reduction with the TI results produced remaining very consistent with the underlying 'ground truth model'. We refrain from a full benchmarking against other methods as they are not designed to handle such high dimensionality. For instance, Slingshot can take several hours even with the dimensionality reachers 50-100 PCs, Monocle and STREAM also require PCA followed by another dimensionality reduction (in the case of STREAM we tried to override these steps resulting in multi-hour runtimes). PAGA and Palantir depend on allowing the inherent subsetting of diffusion components, without which they produce unreasonable answers (e.g. PAGA results in a fully connected cluster graph, Palantir overly fragments the data). As such, we compute the composite accuracy metric for VIA on the full feature sets, but only show some of the TI outputs of PAGA and Palantir when we override any ‘behind-the-scenes’ subsetting of features.

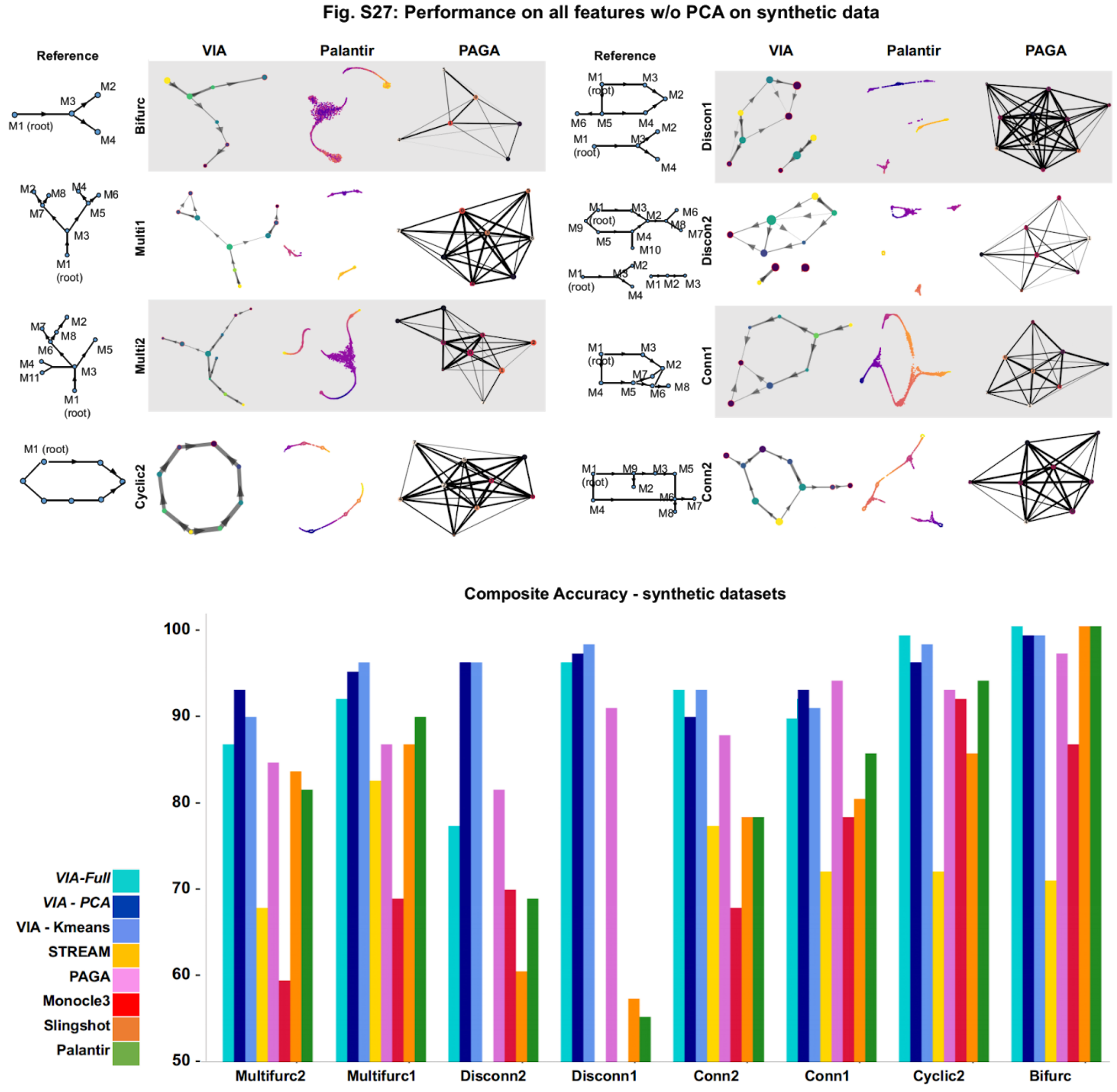


**Fig. S27 TI performance comparison with full feature input using synthetic data (no PCA) (a)** Sample outputs of VIA topology when using all 1000 features without PCA compared to PAGA and Palantir. Although the runtimes for PAGA and Palantir are reasonable, their outputs are distorted due to overriding the PCA and subsetting of diffusion components. **(b)** Composite accuracy scores of VIA when using the full-feature input (VIA-Full), compared to VIA using PCA (VIA-PCA), VIA using Kmeans (and PCA) and all other TI methods which use PCA. We see that VIA can achieve good accuracy even without the PCA step for many of the synthetic datasets.

To see whether VIA TI on full feature inputs can be extended to noisier real data, we extend the analysis to two scRNA-seq datasets (**Fig. S28-S29**), namely pancreatic islet development and human hematopoietic data. In the main text we considered these datasets by following the same pre-processing protocols are their authors: for the hematopoiesis we simply remove low quality cells and genes (resulting in ~15,000 genes across 5780 cells), perform library normalization and log-transformation before applying PCA to the full feature matrix (no subsetting of highly variable genes). We therefore examined the impact on performance across choice of number of nearest neighbors (K) and principal components. For the pancreatic development data the authors performed a similar data filtering and log-normalization (yielding ~15,000 genes across 2531 cells) but then subsetted the first 2000 HVGs before applying PCA. We therefore chose to evaluate the impact of choice of HVGs (300 - 10,000 HVGs) and subsequent number of PCs.

In this case where we are examining the ability of VIA to handle genes as direct input, we consider the choice of number of HVGs provided to the algorithm after filtering and log-normalization, but without any PCA. While it is difficult to quantify the quality of the topology we can visually demonstrate the effect of increasing the number of HVGs for the two datasets by showcasing the outputs alongside each other. We find in general that the hematopoietic dataset yields high quality results when using 500-5000 genes, but that the structure deteriorates if the number of genes retained increases beyond this. This is expected given the number of features would begin to significantly exceed the number of datapoints (cells). Surprisingly, the pancreatic dataset tolerates using up to 10,000 HVGs without sacrificing topology or cell fate prediction, even though the number of cells is only a few thousand.

**Fig. S28: VIA on 500HVG of Hematopoiesis**

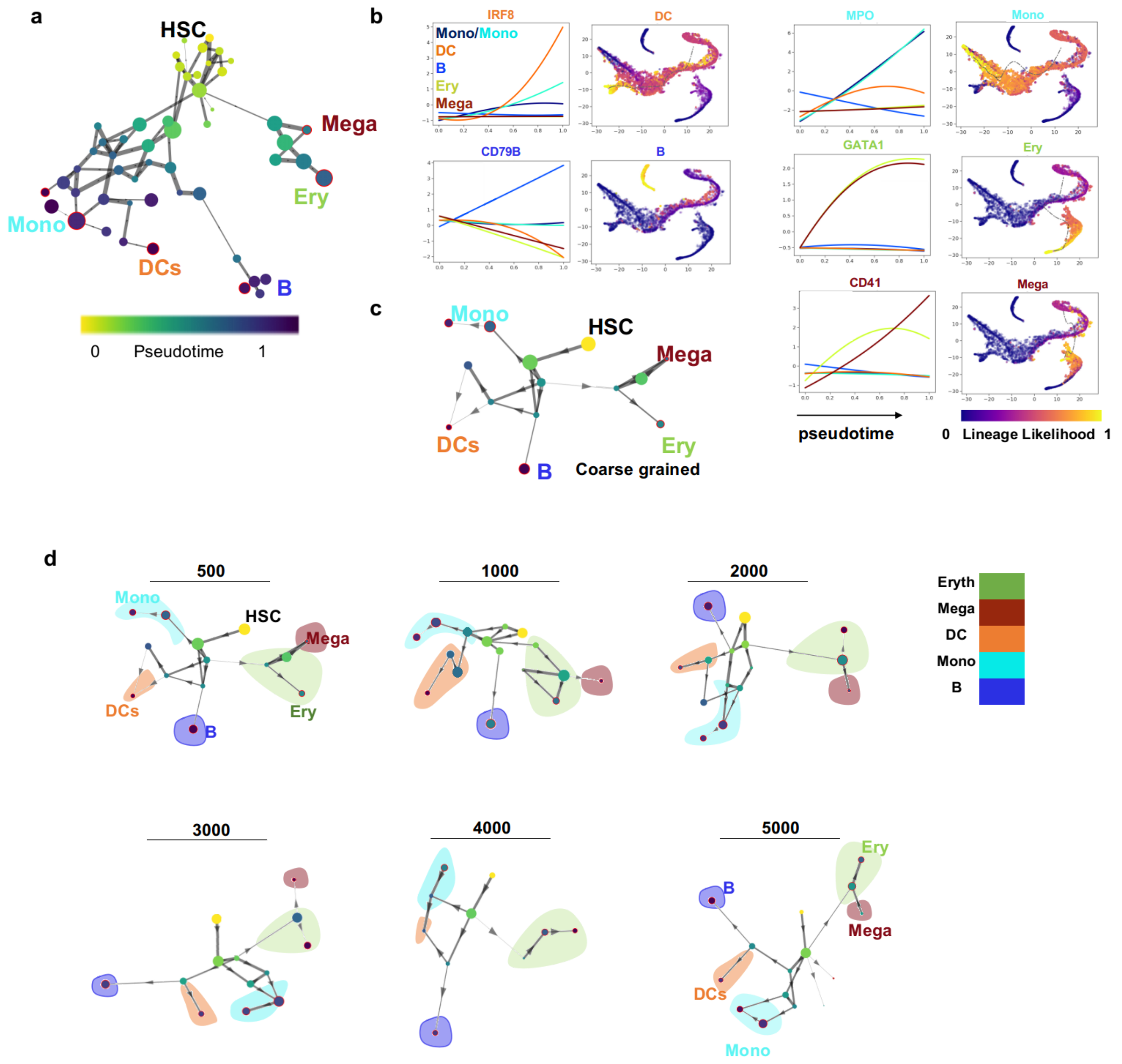


**Fig. S28 Highly variable genes (HVGs) as unprocessed input using the scRNA-seq data of hematopoiesis (no PCA) (a)** VIA graph on 500 HVGs without PCA **(b)** unsupervised plotting of gene trends vs. inferred pseudotimes for predicted lineages. **(c)** Coarse grained VIA graph. **(d)** VIA graph topology for increasing number of HVGs without PCA show that the overall pronged topology is stable with erythroid and megakaryocytic branches on one side of the HSCs, and the monocytes and DCs on the other side.

#### Fig. S29: VIA on HVG of Endocrine genesis

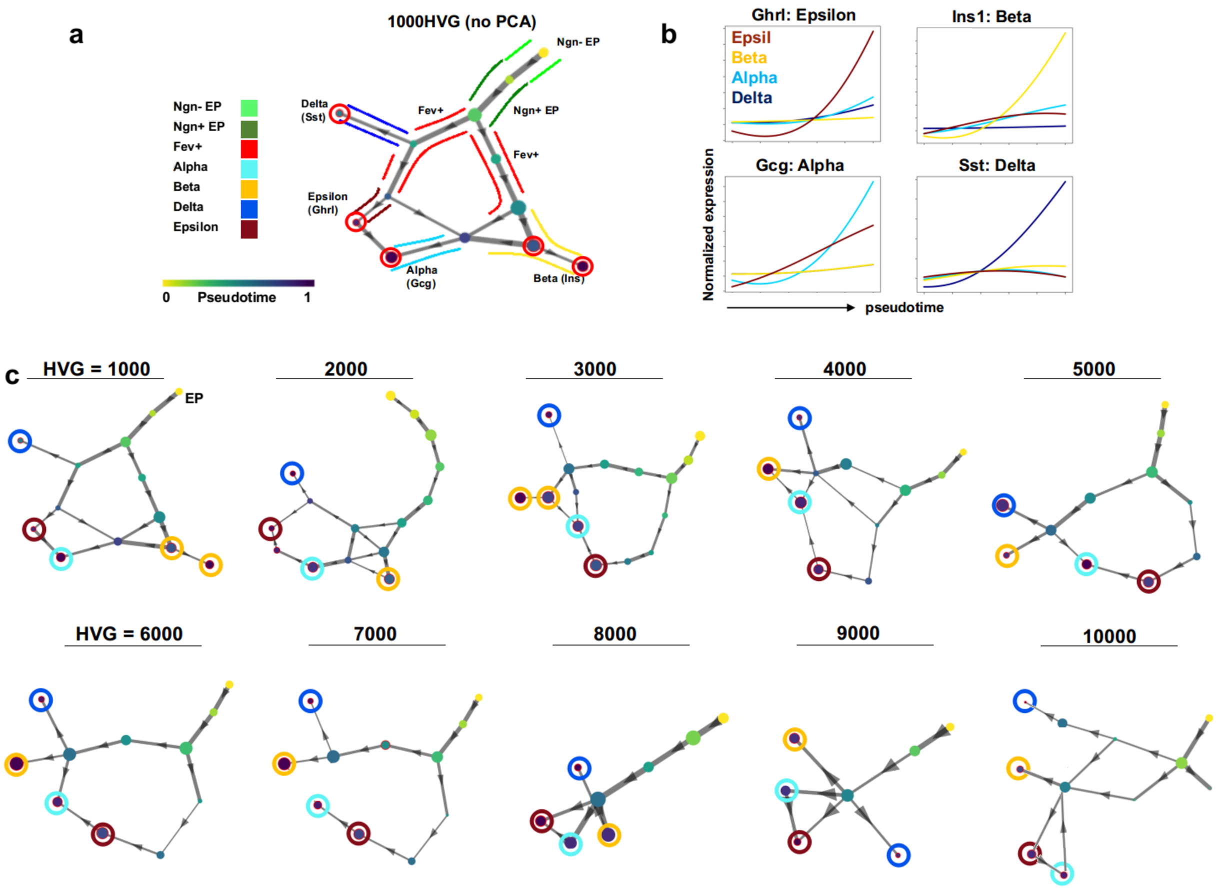


**Fig. S29 Highly variable genes (HVGs) as direct input using the scRNA-seq data of endocrine genesis (no PCA). (a)** Overall graph level topology of VIA, nodes colored by inferred pseudotime and automatically detected terminal nodes circled red. **(b)** Marker gene expression plotted for each lineage shows that each cell fate and pathway uniquely leads to the islet fate associated with that marker gene. **(c)** Topology of endocrine dataset for different number of HVGs, continues to represent the overall progression from EPs to bifurcation of *Fev+* cells that lead to the different islets.

### 11. Supplementary Note 6: VIA performance with different clustering method

A key step in VIA is to reduce the single-cell graph into a cluster-graph using the PARC's community detection algorithm. We assessed the performance of VIA when substituting PARC with another clustering method (in this case we chose Kmeans as it is representative of extremely simple but fast clustering) to show that VIA's teleporting-random walks still traverse and uncover the expected trajectories and their associated lineage-cell-fates.

The analysis shows two key characteristics, the first is that VIA's performance is not limited to using PARC in the clustering step (**Fig. S30a**), and the second is that given the continuous nature of the underlying synthetic data (**Fig. S30b**), VIA's lazy-teleporting random walks still work well even when the data is not inherently highly clustered. We first tested performance on the synthetic datasets and compared the composite metric scores of VIA with PARC and with Kmeans. The composite score allows us to quantify accuracy of topology, pseudotime and F1-cell fate prediction for all synthetic datasets across various topologies and demonstrate that VIA using Kmeans yielded competitive results **(Fig. S30a).**

Following this, we also show that the expected gene trends and progression of cells can be recovered on real data (scRNA-seq hematopoiesis and endocrine genesis). In **Fig. S31-S32**, we not only show that the topology uncovered remains true to the known biological progression towards committed lineages (as verified through reference to literature of the hematopoietic and endocrine genesis processes), but also that the gene expression trends of known marker genes plotted versus the inferred pseudotime for each of the predicted lineages, reflect the expected upregulation. We set 20 as the number of clusters, however, at n=10 clusters, the megakaryocyte cells are absorbed into the larger erythroid lineage, and the dendritic cells are merged together or at times even absorbed into the monocytic lineage).

Thus, a drawback of using Kmeans is that the choice of number of clusters needs to be sensible to capture the desired resolution. In a classical clustering scenario, this might be more problematic as the choice of number of clusters can incur fragmentation of cell types in order to uncover a smaller population, thus generating 'false clusters', or merging of dissimilar cell types if the number of clusters is set to be too low.

To avoid the issue of setting the number of clusters, methods like PARC uncover the underlying granularity in a data-driven manner (without requiring a predetermined number of clusters). In PARC, the underlying graph pruning allows it to automatically detect rarer cell types. However, for the datasets considered, we found that setting K to a sensible number that captures populations of various abundances along the trajectory, still allows the relevant neighborhood information to be subsequently propagated in the random walks. These results ultimately allow the user more flexibility in their choice of how to group or cluster cells or potentially even use prior knowledge of cell types to guide the inferred trajectory.

#### Fig. S30: VIA robust to choice of clustering method

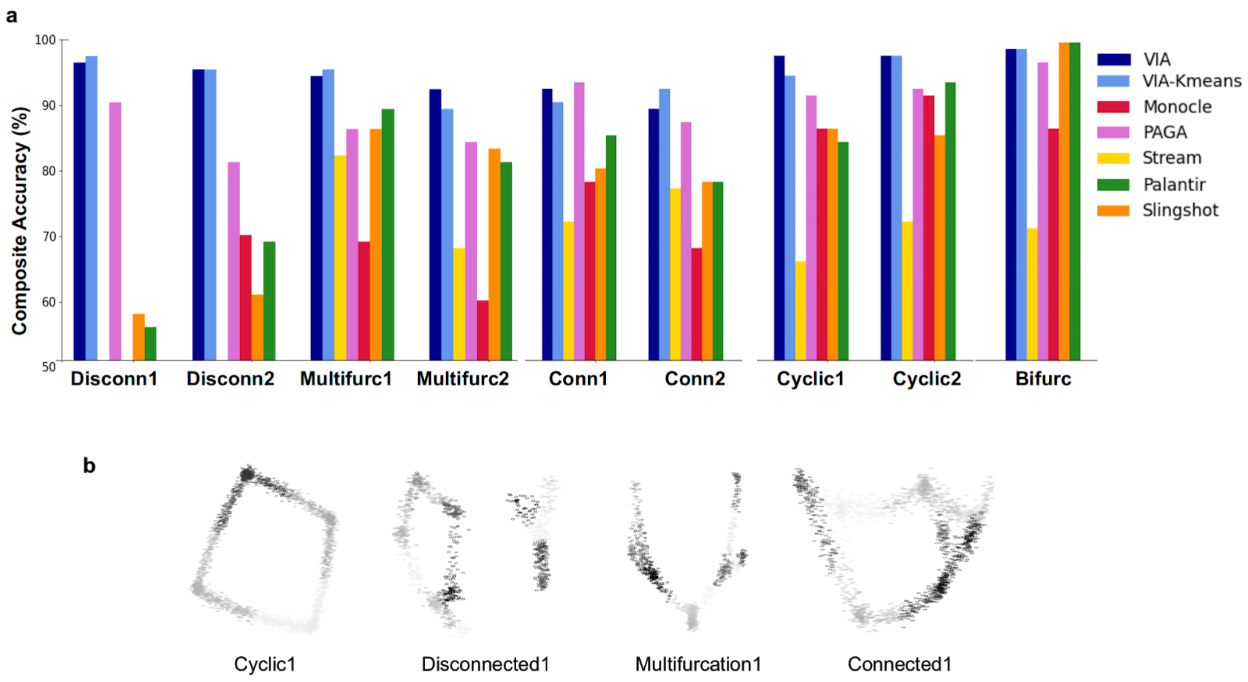


**Fig. S30 VIA with Kmeans clustering in lieu of PARC in the clustering step of VIA for synthetic data maintains accuracy of TI (a)** VIA with Kmeans (VIA-Kmeans) and the original VIA, i.e. VIA with PARC, (VIA) are comparable in terms of performance on synthetic datasets, with both approaches maintaining a competitive advantage over other methods, particularly for multifurcations and disconnected and connected topologies **(b)** Synthetic datasets plotted with first two principal components show the underlying data is continuous and not composed of discrete clusters.

#### Fig. S31: VIA using KMeans on Pancreatic Islets

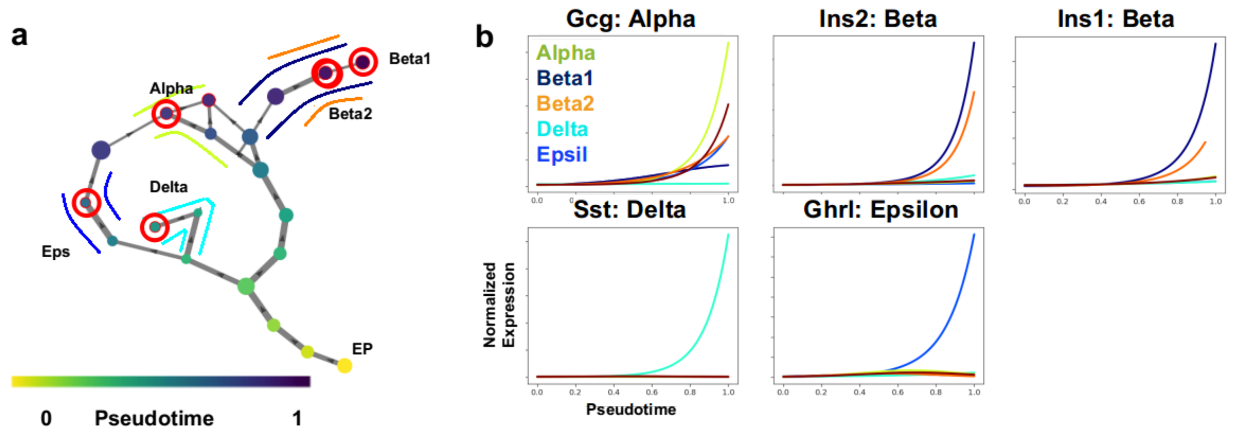


**Fig. S31 VIA with Kmeans clustering in lieu of PARC for endocrine genesis data (a)** At 20 clusters, VIA automatically detects the 4 islets as terminal states (red circles). The topology captures the progression of lineage commitment arising after progressing from *Ngn*-low to -high EPs, followed by intermediate *Fev*+ cells. **(b)** Unsupervised gene expression vs. pseudotime of lineage markers show the unique elevation of specific genes with minimal cross-talk from unrelated lineages. The two Beta subtypes are detected, with the Beta-2 subtype expressing *Ins2* but not *Ins1*.

**Fig. S32: VIA using KMeans on scRNA-seq Hematopoiesis**

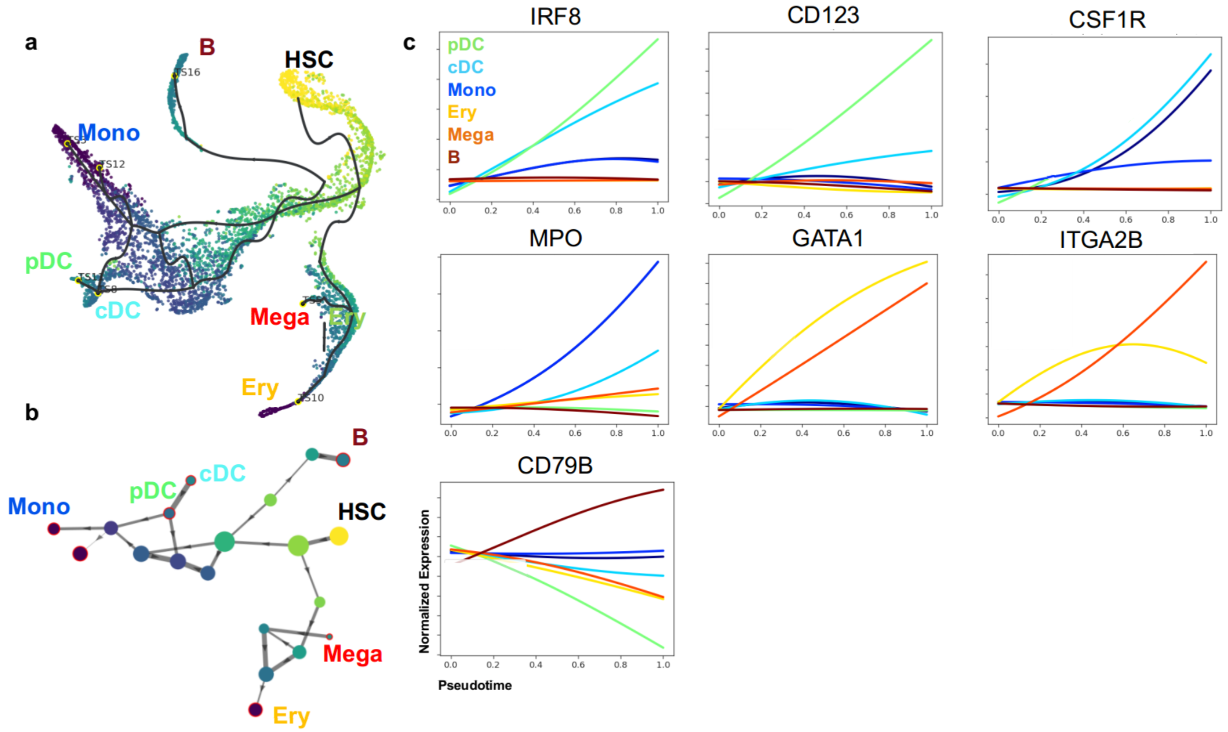


**Fig. S32 VIA with Kmeans clustering in lieu of PARC for scRNA-seq Hematopoiesis (a)** At 20 clusters, VIA automatically detects the 6 lineages specific to the pDC, cDC, monocyte, megakaryocyte, B cell and erythroid cell fates. **(b)** Unsupervised gene expression vs. pseudotime of lineage markers show the unique elevation of specific genes with minimal cross-talk from unrelated lineages. Notably *ITGA2B* (*CD41*) is elevated in the Megakaryocyte lineage and *CD123* is elevated in the DC lineage representing the pDCs, and *CSF1R* is elevated in the cDC lineage.

### 12. Supplementary Note 7: VIA structural connections supported by known transitions in organogenesis literature

VIA's graph layout provides a coarse-grained interpretation of the developmental landscape. Although it remains challenging to capture the simultaneously developing trajectories and cross-talk in a single snapshot, VIA allows lineages to be explored individually (examples are shown in **Supplementary Fig. S11c**) and shows trajectories in agreement with those by Cao et al[17]. In VIA, **the stem branch** contains epithelial cells originating in the ectoderm (such as the epidermis, nose and mouth) as well as those originating in the endoderm (such as the foregut/hindgut)[19]. The epithelial branch also has the largest percentage concentration of E9.5 cells by major trajectory and is therefore designated "root" status. This epithelial stem has two main branches, (1) the mesenchymal cells (derived from the mesoderm) and (2) the neuronal and neural crest cells formed during neurulation when the ectoderm folds inwards[20].

The **mesenchymal cells** arise from the mesoderm which is formed by interactions between the ectoderm and endoderm. The connectivity observed between the epithelial and mesenchymal cells could be supported by epithelial-to-mesenchymal transitions known to occur in the developing embryo[23] when tissues are constructed, while the reverse process (i.e. mesenchymal-to-epithelial) can similarly generate secondary epithelial cells[34,21,22]. Dong et al.[35] even describe a hybrid epithelial-mesenchymal state in a separate study on mouse organogenesis. For instance, signaling in the gut epithelium at E9.5 induces proliferation of the mesenchymal gut tissues and epithelium between E9.5 and E13.5[19,24] In VIA, the junction between epithelial and mesenchymal groups is populated by early mesenchymal cells predominantly from E9.5-E10.5, directly followed by clusters containing limb, chondrocyte and osteoblast cells which are mainly E10.5-E11.5. Myocyte cells from E12.5-E13.5 reside at the leaves.
The **neural-notochord branch** connected to the stem-branch (epithelial-ectoderm cells) contains the neural-tube/notochord cells. During neurulation, the ectoderm folds inwards and forms the neural tube which gives rise to the central nervous system[20]. During neurulation the ectoderm also creates the neural crest tissues (PNS glia, PNS neurons and melanocytes). At the neural tube and epithelial branch junction of VIA, we find clusters largely made of E9.5 and E10.5 dorsal neural tube cells and notochord plate cells. Progressing towards the leaves of this branch, we find excitatory neurons from E12.5-E13.5 stages.

The **Neural crest branch:** Cao et al.[17] identify 3 stand alone neural crest groups that in their UMAP-Monocle3 analysis appear disconnected from each other and other groups. However, it is known that neural crest cells arise at the margins of the neural tube during neurulation, after which they can migrate to many different locations within the embryo[20]. VIA places the cells belonging to the PNS neural crest cells and the PNS glial neural crest cells on a side branch, at the periphery of the major group associated with the neural tube/notochord cell types. The Schwann cell precursors (SCP) making up the glial neural crest super-group are connected to the radial glial cells in the neural tube/notochord branch as well as to the PNS neurons. The Schwann cells together with the PNS neurons form the myelin sheath.

**Other major lineages:** Cao[17] finds a handful of other dispersed cell groups. VIA consistently places these in neighborhoods relevant to their development. In fact, when changing the UMAP parameters, we find that the connectivity (or lack thereof) between the 10 super-groups is in flux, inevitably impacting the

Monocle3 interpretation. We find that changing similar parameters in VIA (such as K number of neighbors, and input PCs) does not perturb fundamental linkages between major groups **(see Fig. S10-14).** The **hepatocyte cells** are polarized epithelial cells originating from the endoderm[30] and hence connected by an edge to the Epithelial stem-branch which includes endoderm derived fore/hind gut epithelial cells. The **hematopoietic cells** and the **endothelial cells** are both derived from the hemangioblast mesodermal cells and both connected to the mesenchymal branch[31] The cluster connecting the mesenchymal branch to the hematopoietic one consists of stromal cells which mechanically support differentiating hematopoietic cells[36]**.**

#### 13. Table S6 FACED image extracted biophysical features and the corresponding definitions[32,33]

| | Feature | Symbol | Equation |
|---|---|---|---|
| Bulk | Area | A | $L_{pix}^{2} \cdot N_{pix}$ |
| | Volume | V | $\frac{4}{3}\pi \cdot \left(\frac{L_{minor}}{2}\right)^{2} \cdot \left(\frac{L_{major}}{2}\right)$ |
| | Circularity | | $4\pi A/P$ |
| | Eccentricity | | $\frac{L_{ellip}}{L_{major}}$ |
| | Aspect Ratio | | $\frac{L_{minor}}{L_{major}}$ |
| | Orientation | | $\theta_{major}$ |
| | Dry Mass | $MD_{total}$ | $\frac{\lambda}{2\pi\alpha}\iint_{A} MD(x,y)\,dxdy$ |
| Global | Dry Mass Density | $\acute{DMD}$ | $\frac{\iint_{A} DMD(x,y)\,dxdy}{N_{pix}}$ |
| | Dry Mass Variance | $\sigma_{DMD}^{2}$ | $\iint_{A} \left(DMD(x,y)-\acute{DMD}\right)^{2} dxdy / \left(N_{pix}-1\right)$ |
| | Dry Mass Skewness | | $\frac{\iint_{A} \left(DMD(x,y)-\acute{DMD}\right)^{3} dxdy / N_{pix}}{\sigma_{DMD}^{3}}$ |
| | Peak Phase | | $max\{MD(x,y)\}$ |
| | Phase Variance | | $\frac{\iint_{A} \left(MD(x,y)-\acute{MD}\right)^{2} dxdy}{N_{pix}-1}$ |
| | Phase Skewness | | $\frac{\iint_{A} \left(MD(x,y)-\acute{MD}\right)^{3} dxdy / N_{pix}}{\sigma_{MDstd}^{3}}$ |
| | Phase Kurtosis | | $\frac{\iint_{A} \left(MD(x,y)-\acute{MD}\right)^{4} dxdy / N_{pix}}{\sigma_{MDstd}^{4}}$ |
| | Phase Range | | $max\{MD(x,y)\}-min\{MD(x,y)\}$ |
| | Phase Minimum | | $min\{MD(x,y)\}$ |
| | Phase Centroid Displacement | | $\sqrt{(x_{DMD,cen}-x_{cen})^{2}+(y_{DMD,cen}-y_{cen})^{2}} \cdot L_{pix}$ |

| | | | |
|---|---|---|---|
| **Local** | Mean Phase Arrangement | | $\frac{\iint_A MD(r,\theta)\, r\, d\, rd\theta}{\iint_A MD(r,\theta)\, d\, rd\theta}$ |
| | Phase Arrangement Variance | $\sigma_{MDarr}^{2}$ | $\frac{\iint_A (MD(r,\theta)\, r)^2\, d\, rd\theta}{\iint_A MD(r,\theta)\, d\, rd\theta}$ |
| | Phase Arrangement Skewness | | $\frac{\iint_A (MD(r,\theta)\cdot r)^3\, d\, rd\theta}{\sigma_{PGarr}^{2}\cdot \iint_A MD(r,\theta)\, d\, rd\theta}$ |
| | Phase Orientation Variance | $\sigma_{MDang}^{2}$ | $\frac{\int_0^{\infty} (\widetilde{MD}(\omega)\cdot\omega)^2\, d\,\omega}{\int_0^{\infty} \widetilde{MD}(\omega)\, d\,\omega}$ |
| | Phase Orientation kurtosis | | $\frac{\int_0^{\infty} (\widetilde{MD}(\omega)\cdot\omega)^4\, d\,\omega}{\sigma_{PGang}^{2}\cdot \int_0^{\infty} \widetilde{MD}(\omega)\, d\,\omega}$ |
| | Phase STD Mean | $\acute{MD}_{STD}$ | $\frac{\iint_A MD_{STD}(x,y)\, d\, xdy}{N_{pix}}$ |
| | Phase STD Variance | $\sigma_{MDstd}^{2}$ | $\frac{\iint_A (MD_{STD}(x,y) - \acute{MD}_{STD})^2\, d\, xdy}{N_{pix}-1}$ |
| | Phase STD Skewness | | $\frac{\iint_A (MD_{STD}(x,y) - \acute{MD}_{STD})^3\, d\, xdy / N_{pix}}{\sigma_{MDstd}^{3}}$ |
| | Phase STD Kurtosis | | $\frac{\iint_A (MD_{STD}(x,y) - \acute{MD}_{STD})^4\, d\, xdy / N_{pix}}{\sigma_{MDstd}^{4}}$ |
| | Phase STD Centroid Displacement | | $\sqrt{(x_{MDSTD,cen} - x_{cen})^2 + (y_{MDSTD,cen} - y_{cen})^2}\cdot L_{pix}$ |
| | Phase STD Radial Distribution | | $\frac{\iint_A r\cdot MD_{STD}(r,\theta)\, d\, rd\theta}{\iint_A MD_{STD}(r,\theta)\, d\, rd\theta}$ |
| | Phase Entropy Mean | $\acute{MD}_{ent}$ | $\frac{\iint_A MD_{ent}(x,y)\, d\, xdy}{N_{pix}}$ |

| | | | |
|---|---|---|---|
| | Phase Entropy Variance | $\sigma_{MDent}{}^{2}$ | $$\frac{\iint_A \left(MD_{ent}(x,y)-\acute{MD}_{ent}\right)^2 d\,xdy}{N_{pix}-1}$$ |
| | Phase Entropy Skewness | | $$\frac{\iint_A \left(MD_{ent}(x,y)-\acute{MD}_{ent}\right)^3 d\,xdy/N_{pix}}{\sigma_{MDent}{}^{3}}$$ |
| | Phase Entropy Kurtosis | | $$\frac{\iint_A \left(MD_{ent}(x,y)-\acute{MD}_{ent}\right)^4 d\,xdy/N_{pix}}{\sigma_{MDent}{}^{4}}$$ |
| | Phase Entropy Centroid Displacement | | $$\sqrt{(x_{MDent,cen}-x_{cen})^2+(y_{MDent,cen}-y_{cen})^2}\cdot L_{pix}$$ |
| | Phase Entropy Radial Distribution | | $$\frac{\iint_A r\cdot MD_{ent}(r,\theta)\,d\,rd\theta}{\iint_A MD_{ent}(r,\theta)\,d\,rd\theta}$$ |
| | Phase Fiber Centroid Displacement | | $$\sqrt{(x_{MDfiber,cen}-x_{cen})^2+(y_{MDfiber,cen}-y_{cen})^2}\cdot L_{pix}$$ |
| | Phase Fiber Radial Distribution | | $$\frac{\iint_A r\cdot MD_{fiber}(r,\theta)\,d\,rd\theta}{\iint_A MD_{fiber}(r,\theta)\,d\,rd\theta}$$ |
| | Phase Fiber Pixel > Upper Percentile | | $$\frac{Number\ of\ pixels \in MD_{fiber}(x,y) > 75\,th\ percentile}{N_{pix}}$$ |
| | Phase Fiber Pixel > Median | | $$\frac{Number\ of\ pixels \in MD_{fiber}(x,y) > median}{N_{pix}}$$ |

**Table S7 List of variables (descriptions and definitions) used in the Equations (Table S6)**

| Variable | Description | Equation |
|---|---|---|
| $C$ | Contour of binary mask | |
| $CM$ | Cell mask function | $CM(x,y)=\begin{cases}1 \; if\ inside\ cell\\ 0 \;\; otherwise\end{cases}$ |
| $DMD$ | Dry mass density map | $DMD(x,y)=\frac{\lambda \cdot MD(x,y)}{2\pi\alpha \cdot h(x,y)}$ |
| $h$ | Cell height map | $h(x,y)=\sqrt{\left(\frac{L_{minor}+L_{major}}{2}\right)^2-\left((x-x_{cen})^2+(y-y_{cen})^2\right)}$ |
| $L_{ellip}$ | Distance between foci of ellipse | |
| $L_{major}$ | Major axis length | |
| $L_{minor}$ | Minor axis length | |
| $L_{pix}$ | Physical length of one pixel | |
| $MD$ | Mass density map | $MD(x,y)$ |
| $MD(\theta)$ | MD projected to polar angle | |
| $\widetilde{MD}(\omega)$ | MD in angular frequency domain | $\widetilde{MD}(\omega)=F(MD(\theta))$ |
| $MD'_{STD,ker}(x,y)$ | Mean value of QPI within STD filter kernel | $\frac{\int_{x-w_{STD}/2}^{x+w_{STD}/2}\int_{y-w_{STD}/2}^{y+w_{STD}/2} MD(u,v)\,dvdu}{w_{STD}^2}$ |
| $MD_{STD}(x,y)$ | QPI STD map | $\int_{x-w_{STD}/2}^{x+w_{STD}/2}\int_{y-w_{STD}/2}^{y+w_{STD}/2}\sqrt{\frac{(MD(u,v)-MD'_{STD,ker}(x,y))^2}{w_{STD}^2}}\,dvdu$ |
| $MD_{cubic}(x,y)$ | Cubic polynomial surface fit of MD | |
| $MD_{ent}(x,y)$ | Entropy filtered MD | $\sum_{k=0}^{255} p_{MD,k}\cdot\log_2 p_{MD,k}$ |
| $MD_{fiber}(x,y)$ | Fiber texture enhanced MD | $FF(MD(x,y))$ , ref. [2] |
| $N_{pix}$ | Pixel number in cell mask | $\iint CM(x,y)\,dA$ |
| $P$ | Perimeter | $\oint_C \sqrt{\left(\left(\frac{dx}{d\theta}\right)^2+\left(\frac{dy}{d\theta}\right)^2\right)}\,d\theta$ |
| $p_{MD,k}(x,y)$ | Normalized histogram counts within kernel of MD | $p_{MD,k}(x,y)=\frac{number\ of\ pixels \in kernel(w_{ent})\ with\ MD=k}{Total\ number\ of\ pixels \in kernel}$ |

| | | |
|---|---|---|
| $r, \theta$ | Polar coordinates centered at cell centroid | |
| $w_{ent}$ | Kernel size of entropy filter | |
| $w_{STD}$ | Kernel size of STD filter | |
| $x, y$ | Cartesian coordinates | |
| $x_{cen}$<br>$y_{cen}$ | Coordinates of cell centroid | $x_{cen} = \frac{\iint_A x \cdot CM(x,y)\,dxdy}{N_{pix}}$<br>$y_{cen} = \frac{\iint_A y \cdot CM(x,y)\,dxdy}{N_{pix}}$ |
| $x_{MD,cen}$<br>$y_{MD,cen}$ | Coordinates of MD weighted cell centroid | $x_{MD,cen} = \frac{\iint_A x \cdot MD(x,y)\,dxdy}{N_{pix}}$<br>$y_{MD,cen} = \frac{\iint_A y \cdot MD(x,y)\,dxdy}{N_{pix}}$ |
| $x_{DMD,cen}$<br>$y_{DMD,cen}$ | Coordinates of DMD weighted cell centroid | $x_{MDDcen} = \frac{\iint_A x \cdot DMD(x,y)\,dxdy}{N_{pix}}$<br>$y_{MDDcen} = \frac{\iint_A y \cdot DMD(x,y)\,dxdy}{N_{pix}}$ |
| $x_{MDent,cen}$<br>$y_{MDent,cen}$ | Coordinates of entropy filtered MD weighted cell centroid | $x_{MDent,cen} = \frac{\iint_A x \cdot MD_{ent}(x,y)\,dxdy}{N_{pix}}$<br>$y_{MDent,cen} = \frac{\iint_A y \cdot MD_{ent}(x,y)\,dxdy}{N_{pix}}$ |
| $x_{MDfiber,cen}$<br>$y_{MDfiber,cen}$ | Coordinates of fiber enhanced MD weighted cell centroid | $x_{MDfiber,cen} = \frac{\iint_A x \cdot MD_{fiber}(x,y)\,dxdy}{N_{pix}}$<br>$y_{MDfiber,cen} = \frac{\iint_A y \cdot MD_{fiber}(x,y)\,dxdy}{N_{pix}}$ |
| $x_{MDSTD,cen}$<br>$y_{MDSTD,cen}$ | Coordinates of STD filtered MD weighted cell centroid | $x_{MDSTD,cen} = \frac{\iint_A x \cdot MD_{STD}(x,y)\,dxdy}{N_{pix}}$<br>$y_{MDSTD,cen} = \frac{\iint_A y \cdot MD_{STD}(x,y)\,dxdy}{N_{pix}}$ |
| $\alpha$ | Specific refractive increment | $0.19\,ml/g$ (ref. [3]) |
| $\theta_{major}$ | Angle between major axis and x-axis | |
| $F$ | Fourier transform | |
